# Coordinated activation of both TGFβ and BMP canonical pathways regulates autophagy and tissue regeneration in acetaminophen induced liver injury

**DOI:** 10.1101/2021.07.23.453185

**Authors:** Athanasios Stavropoulos, Georgios Divolis, Maria Manioudaki, Ariana Gavriil, Ismini Kloukina, Despina N. Perrea, Alexandros Sountoulidis, Ethan Ford, Athanasia Doulou, Anastasia Apostolidou, Elena Katsantoni, Olli Ritvos, Georgios Germanidis, Maria Xilouri, Paschalis Sideras

**Author notes:** To whom correspondence should be addressed: Paschalis Sideras, Center of Clinical, Experimental Surgery and Translational Research, Biomedical Research Foundation of the Academy of Athens, Athens, Greece, E- mail.

## Abstract

Transforming Growth Factor-βs (TGFβs)/Activins and Bone Morphogenetic Proteins (BMPs) have been implicated in numerous aspects of hepatic pathophysiology. However, the way by which hepatocytes integrate and decode the interplay between the TGFβ/Activin and BMP branches in health and disease is still not fully understood. To address this, TGFβ/BMP Smad-responsive double transgenic reporter mice were generated and utilized to map patterns of TGFβ- and/or BMP-pathway activation during acetaminophen-induced liver injury. TGFβ signaling was blocked either pharmacologically or by Smad7 over-expression and the transcriptomes of canonical TGFβ- and/or BMP4-treated hepatospheres and Smad7-treated livers were analyzed to highlight TGFβ-superfamily-regulated pathways and processes. Acetaminophen administration led to dynamically evolving, stage- and context-specific, patterns of hepatic TGFβ/Activin and BMP-reporter expression. TGFβ-superfamily signaling was activated in an autophagy prone zone at the borders between healthy and injured tissue. Inhibition of TGFβ-superfamily signaling attenuated autophagy, exacerbated liver histopathology, and finally led to accelerated tissue-recovery. Hallmarks of this process were the paraptosis-like cell death and the attenuation of immune and reparatory cell responses. Transcriptomic analysis highlighted autophagy as a prominent TGFβ1- and BMP4-regulated process and recognized *Trp53inp2* as the top TGFβ-superfamily-regulated autophagy-related gene. Collectively, these findings implicate the coordinated activation of both canonical TGFβ-superfamily signalling branches in balancing autophagic response and tissue-reparatory and -regenerative processes upon acetaminophen-induced hepatotoxicity, highlighting opportunities and putative risks associated with their targeting for treatment of hepatic diseases.

## Introduction

The TGFβ-superfamily signaling system comprises approximately 33 members most of which are gathered in two major groups, the TGFβ/Activins and the Bone Morphogenetic Proteins (BMPs) (1). These ligands signal to the nucleus via phosphorylation of dedicated effectors, the Smad proteins. TGFβs and Activins induce phosphorylation and activation of Smads2/3, while BMPs induce phosphorylation and activation of Smads1/5/8 (2), thus establishing two signaling branches. Activated Smads form complexes with Smad4 and translocate to the nucleus where they regulate gene expression. Smad6 and Smad7 function as feedback negative regulators of the system. In parallel with Smad-dependent, “canonical” pathways, TGFβ-superfamily ligands can also activate Smad-independent, “non-canonical” pathways (3).

Several ligands of both TGFβ and BMP signaling branches have been implicated in various aspects of liver pathophysiology. TGFβ was found increased in liver-fibrosis models (4–6) and human cirrhosis (7), hepatic cancer (8, 9), alcohol-damage (10), viral hepatitis (11, 12) and was associated with detrimental or protective functions. BMP-signaling has been assigned a key role in regulating hepatic iron-metabolism (13, 14) via hepcidin (15). The role of BMP-signaling in liver-fibrosis is complex. BMP7 exhibits anti-fibrotic (16) and BMP9 pro-fibrotic roles (17). Inhibition of TGFβ-signaling enhances liver-regeneration in models of partial hepatectomy (18, 19), while the effect of BMP-signaling is controversial (17, 20, 21). Acetaminophen (also known as APAP or paracetamol) is a well-known analgesic/antipyretic drug. However, it is the leading cause of acute liver failure and intensive care unit hospitalizations due to chronic drug administration or overdose (22), and liver transplantation in the western world (23). Recent studies have highlighted a negative role of TGFβ-signaling in liver regeneration after APAP-induced liver injury (24, 25).

So far, the only link between the BMP-pathway and APAP pathophysiology is the APAP-induced suppression of BMP-signaling, and subsequently, of hepcidin expression, leading to hepatic iron loading that contributes to hepatotoxicity (26).

To shed light on the rules that govern the balance between the TGFβ/Activin- and BMP-signaling branches and clarify the role of this signaling system in the context of APAP pathophysiology we generated and analyzed reporter mice enabling simultaneous *in-situ* visualization of canonical TGFβ- and/or BMP-signaling. Our studies unveiled a dynamically evolving interplay between these two signaling branches and provided novel insights into their pleiotropic role in liver pathophysiology, particularly in regulating autophagy. Prospectively, these findings will further enhance understanding of the role of the TGFβ-superfamily signaling system in hepatic pathophysiology and aid the development of novel therapeutic interventions.

## Results

### Development and validation of a TGFβ/Activin & BMP double-reporter mouse line

To visualize *in-situ* activation of canonical TGFβ/Activin signaling, we generated reporter animals, utilizing a well-characterized Smad2/3 responsive element (27) to drive expression of mRFP1 (monomeric Red Fluorescent protein 1, hereafter referred to as RFP) (28) (Supplementary Figure 1A). Hepatospheres formed by primary hepatocytes derived from TGFβ-Responsive-Element RFP-expressing animals (TRE-RFP) (29) were treated with increasing levels of TGFβ1, BMP4, TGFβ receptor-selective inhibitors (SB431542 and LY364947), or indicated combinations of the above (Supplementary Figure 1B-C). TGFβ1 administration evoked a dose-depended reporter activation, which was abolished by the TGFβ inhibitors. On the contrary, BMP4 did not alter RFP expression, even at high doses, thus confirming reporter selectivity. To simultaneously monitor activation of both TGFβ/Activin and BMP canonical pathways, TRE-RFP reporter mice were crossed to previously developed Smad1/5/8-responsive, eGFP-expressing, reporter mice herein referred to as BRE- eGFP (30). TGFβ and BMP selectivity of the double-reporter model, herein referred to as TRE-BRE, was validated *in-vivo* using adenovirus-mediated overexpression of Smad3 or Smad1 in the liver. RFP expression was significantly induced upon intravenous injection of Smad3 adenovirus, whereas eGFP expression was specifically augmented by Smad1 adenovirus (Supplementary Figure 1D-E).

Immunohistochemical analysis of tissue sections from various organs of adult TRE- BRE mice demonstrated that the reporters exhibited consistent, tissue- and cell-specific, as well as intriguingly interrelated and overlapping spatiotemporal expression-patterns (Figure 1A). Among the organs analyzed, pancreas displayed the highest fluorescence intensity, followed by liver, kidney, and intestinal tissues. In the liver, reporter expression was visible as early as embryonic day 14.5, the earliest time point analyzed, and underwent dramatic spread between postnatal days 3-7, involving practically almost all hepatocytes. Thereafter, the reporter positive domains exhibited a gradual, age-dependent contraction towards the central-zones (Figure 1B, Supplementary Figure 2A). Independent validation of the reporters and their expression patterns was provided by demonstrating that levels of pSMAD1/5/8, pSMAD2 and pSMAD3 at different stages of liver development followed faithfully the kinetics of reporter expression (Figure 1C-D).

**Figure 1.**
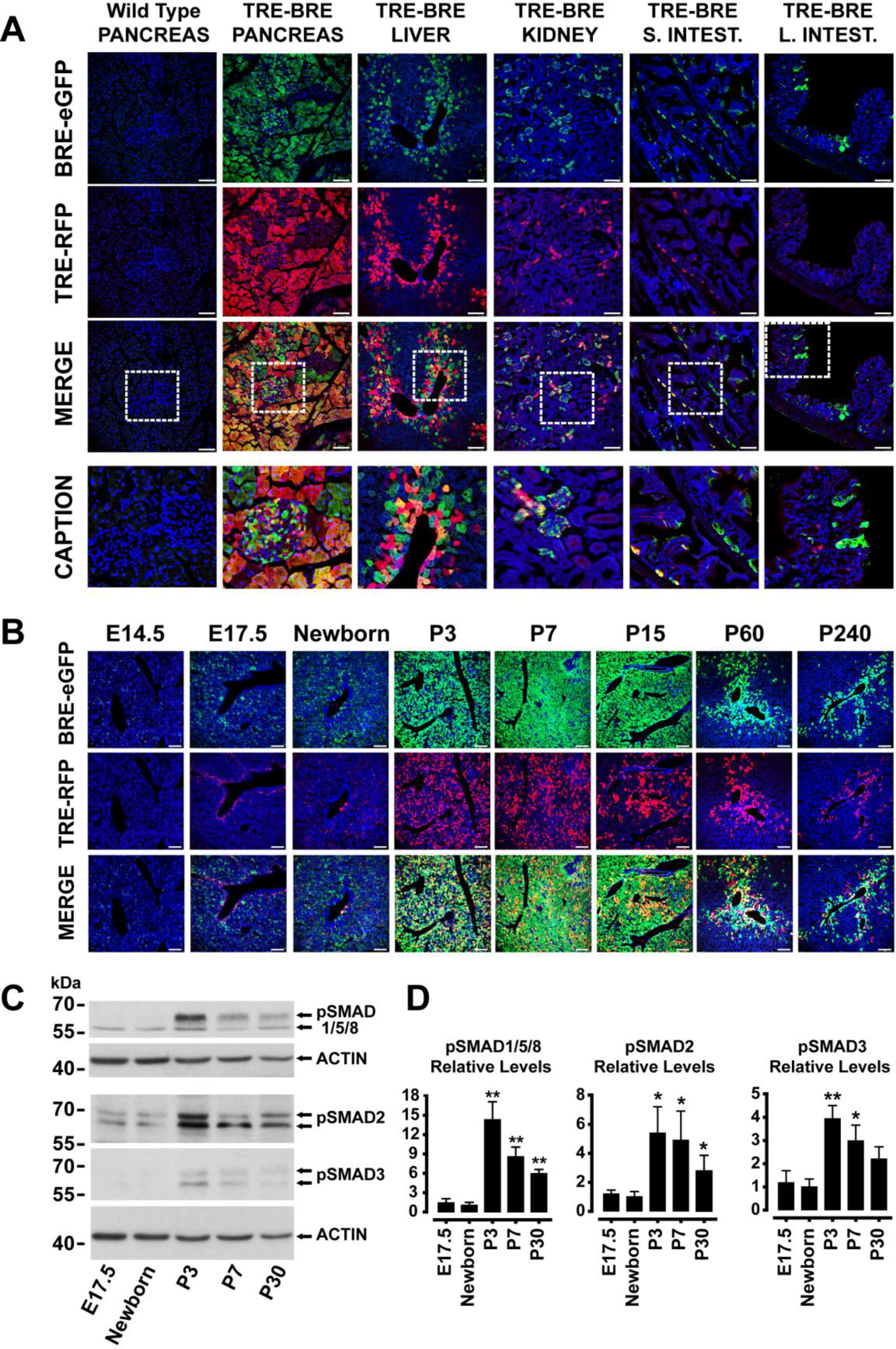
Expression of RFP and eGFP reporters in various organs of adult animals and during hepatic development. (**A**) Representative images of tissue sections from pancreas, liver, kidney, small and large intestine of female TRE-BRE double-reporter or wild-type control animals (pancreas), visualized for eGFP (green), RFP (red), and DAPI (blue). The bottom row represents magnified areas outlined with dashed-lines in the “merged” images directly above (scale-bars, 100μm). (**B**) Representative confocal images of hepatic tissue sections from double-reporter animals at indicated developmental stages (E14.5-P240), stained for eGFP (green), RFP (red), and DAPI (blue), (scale-bars, 100μm). (**C**) Representative western-blots demonstrating similar kinetics for pSMAD1/5/8, pSMAD2, and pSMAD3 levels and expression of the eGFP and RFP reporters during development (β-actin was used as a loading control). (**D**) Quantitative analysis of relative pSMAD/β-actin protein levels normalized to the expression levels of newborn livers. Data are expressed as mean±SEM of 4 independent samples per condition analyzed using one-way analysis of variance with Bonferroni’s post-hoc test.

### Activation of the TGFβ-superfamily signaling system and alterations in autophagy indices upon acetaminophen-induced acute liver injury

To investigate the functional interplay between TGFβ/Activin and BMP signaling in the context of hepatic pathophysiology, we subjected double transgenic TRE-BRE animals to an APAP overdose model, and monitored pathology-driven changes in the spatiotemporal pattern of reporter expression. APAP treatment led to rapid pericentral necrosis (22) and loss of reporter expression at 6h, which was followed by intense re-expression of both reporters in ring-shaped zones surrounding the central-veins, at 24h (Figure S2B). The reporter positive zones were characterized by intense and punctuated expression of p62, Ubiquitin, and LC3, which may be consistent with disturbances in macroautophagy induction or completion at the borders between damaged and non-damaged hepatocytes (Figure 2A). Analysis of the areas corresponding to the reporter positive zones by electron-microscopy (EM) supported this notion by demonstrating accumulation of autophagosomes at different maturation stages, engulfing mainly swollen mitochondria, suggesting active mitophagy (Figure 2B).

**Figure 2.**
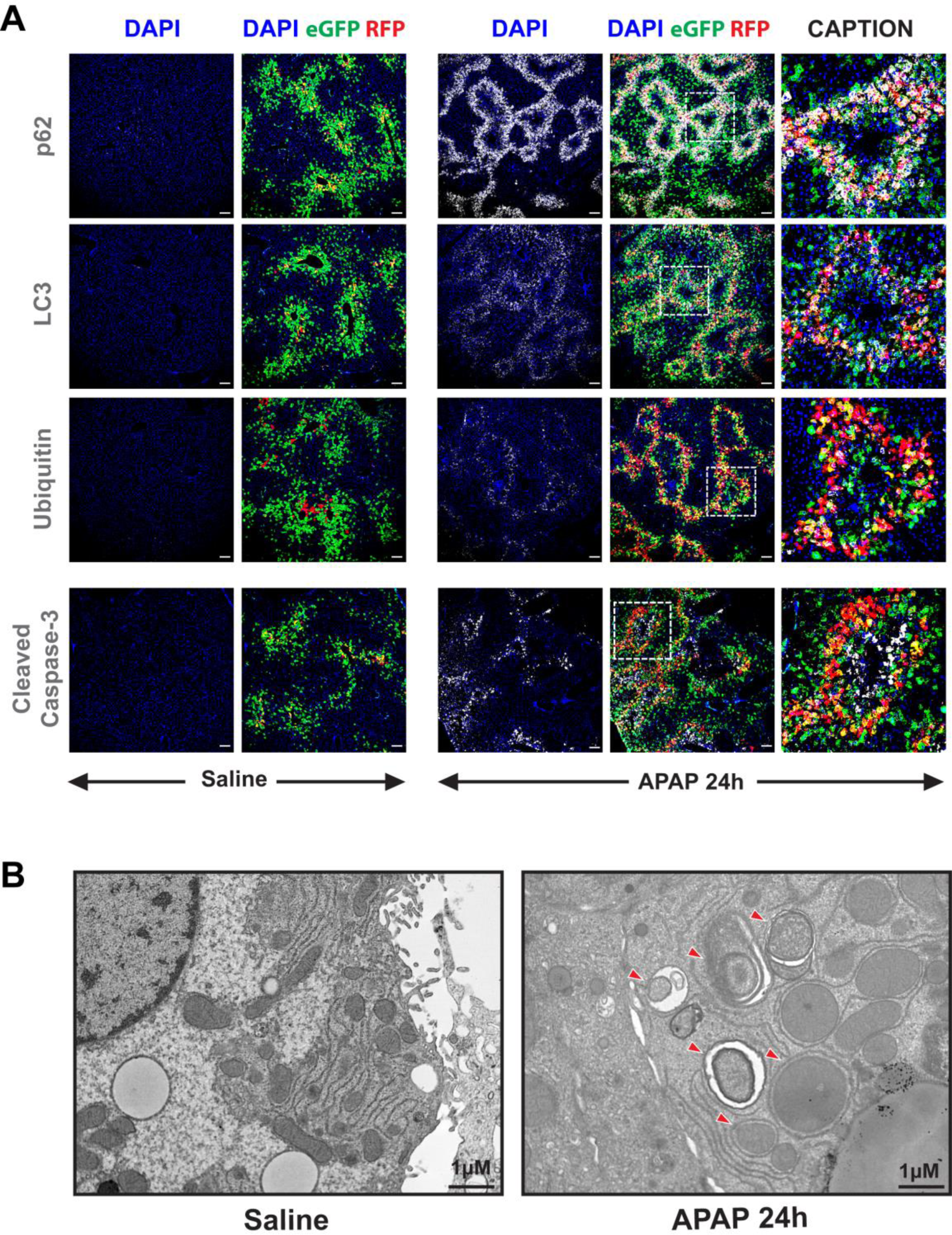
A zone of intense eGFP/RFP reporter co-expression and autophagic activity separates injured from non-injured hepatocytes upon acetaminophen- induced acute liver injury. (**A**) Representative images of hepatic tissue sections from female TRE-BRE reporter animals 24h after APAP or saline administration, visualized for eGFP (green), RFP (red), DAPI (blue) and p62, LC3, ubiquitin or cleaved caspase-3 (white), (scale-bars, 100μm). (**B**) Abundant double-membrane autophagosomes at different maturation stages (indicated by red arrowheads) detected by electron microscopy in the reporter positive zones of APAP-treated livers (scale- bars, 1μm).

### Inhibition of TGFβ-superfamily signaling hinders autophagy, aggravates APAP- induced liver injury but also accelerates tissue recovery

To assess whether activation of canonical TGFβ and/or BMP pathways has a protective or detrimental role in the context of APAP intoxication, Smad7, the endogenous negative regulator of TGFβ/BMP signaling, was ectopically expressed in mouse livers prior to drug administration. Smad7 expression reduced to basal levels APAP-induced phosphorylation of SMAD2 and SMAD1/5/8, at 6h and 24h (Figure 3A-B), delayed clearance of glutathionylated hepatic protein adducts (Figure 3C-D) and interestingly, exacerbated APAP-induced centrilobular necrosis (Figure 3E-F).

**Figure 3.**
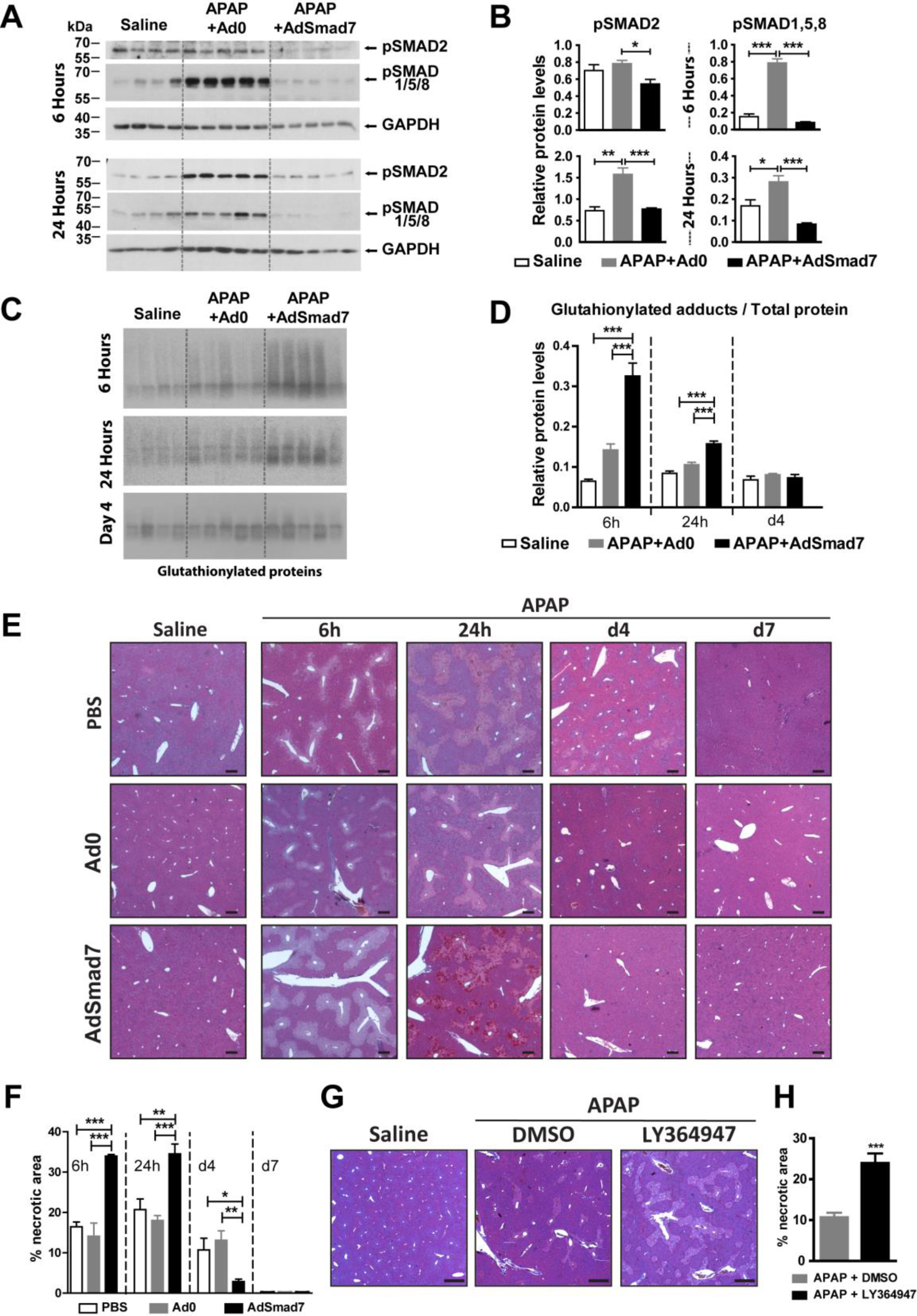
Inhibition of TGFβ-signaling exacerbates acetaminophen-induced hepatic histopathology but accelerates tissue repair. (**A**) Western-blot analysis of pSMAD1/5/8 and pSMAD2 levels. (**B**) Quantitative analysis of pSMADs, relative to GAPDH, in saline or adenovirus plus APAP-treated male animals. (**C**) Representative immunoblots of glutathionylated hepatic proteins in saline or adenovirus plus acetaminophen treated animals, 6h, 24h, and 4days after APAP administration. (**D**) Quantitative analysis of the glutathionylated proteins relative to Ponceau-S stained bands. The data in B and D are expressed as mean±SEM of 4-5 independent samples per group, analyzed using one-way analysis of variance with Bonferroni’s post-hoc test. (**E**) H&E stained tissue sections from male animals injected intravenously with PBS, Ad0 or AdSmad7, treated 16 hours later by intraperitoneal injection of saline or APAP (day-0) and sacrificed at the indicated time-points (scale-bars, 100μm). (**F**) Quantification of liver necrotic areas by ImageJ assisted analysis. Values represent the average of two independent calculations, by two independent observers. Data are expressed as mean±SEM from at least 5 independent samples per group compared at each time point using two-tailed unpaired Student’s t-test. (**G**) H&E staining of tissue sections collected from male animals treated intraperitoneally (i.p.) with saline or APAP followed 1.5h later by i.p. injections of DMSO or LY364947 and sacrificed 24h later (scale-bars, 1mm). (**H**) Quantification of the liver necrotic areas by ImageJ assisted analysis. Data are expressed as mean±SEM from 5 independent samples per group analyzed using two-tailed unpaired Student’s t-test.

Intriguingly, and in agreement with recent findings (24), Smad7 overexpression led to accelerated healing, as exemplified by almost complete tissue restoration four days post-APAP administration instead of the seven days required for the Ad0 treated group (Figure 3E-F). Similar to Smad7 response, prophylactic intraperitoneal administration of the selective TGFβ receptor inhibitor LY364947 significantly aggravated centrilobular necrosis at 24h (Figure 3G-H).

Given the established protective role of autophagy in APAP intoxication (31, 32), we hypothesized that the Smad7-mediated aggravation of histopathology could be due to an effect on hepatocyte autophagy. Thus, we assessed the effect of Smad7 overexpression on the levels of key autophagic proteins of the treated tissues. In agreement with the immunostaining results (Figure 2A), western blot analysis demonstrated that at 6h and 24h post-APAP treatment, levels of p62 and LC3 II proteins that are the hallmarks of the autophagic flux, were substantially elevated in the APAP+Ad0-treated mice compared to the saline controls (Figure 4A-B). In contrast, compared to APAP+Ad0 treated group, protein extracts from APAP+AdSmad7-treated livers, contained lower LC3 II protein levels at both 6 and 24 h (Figure 4A-B) and reduced levels of ATG9 and ATG7 proteins at 6h (Figure 5A) and p62, ATG5 and ATG7 protein levels at 24h (Figure 4A-B and Figure 5A-B). These findings indicate that Smad7 overexpression suppresses autophagic activity most likely via obstruction of cargo-protein recycling at an early stage of autophagosome formation.

**Figure 4.**
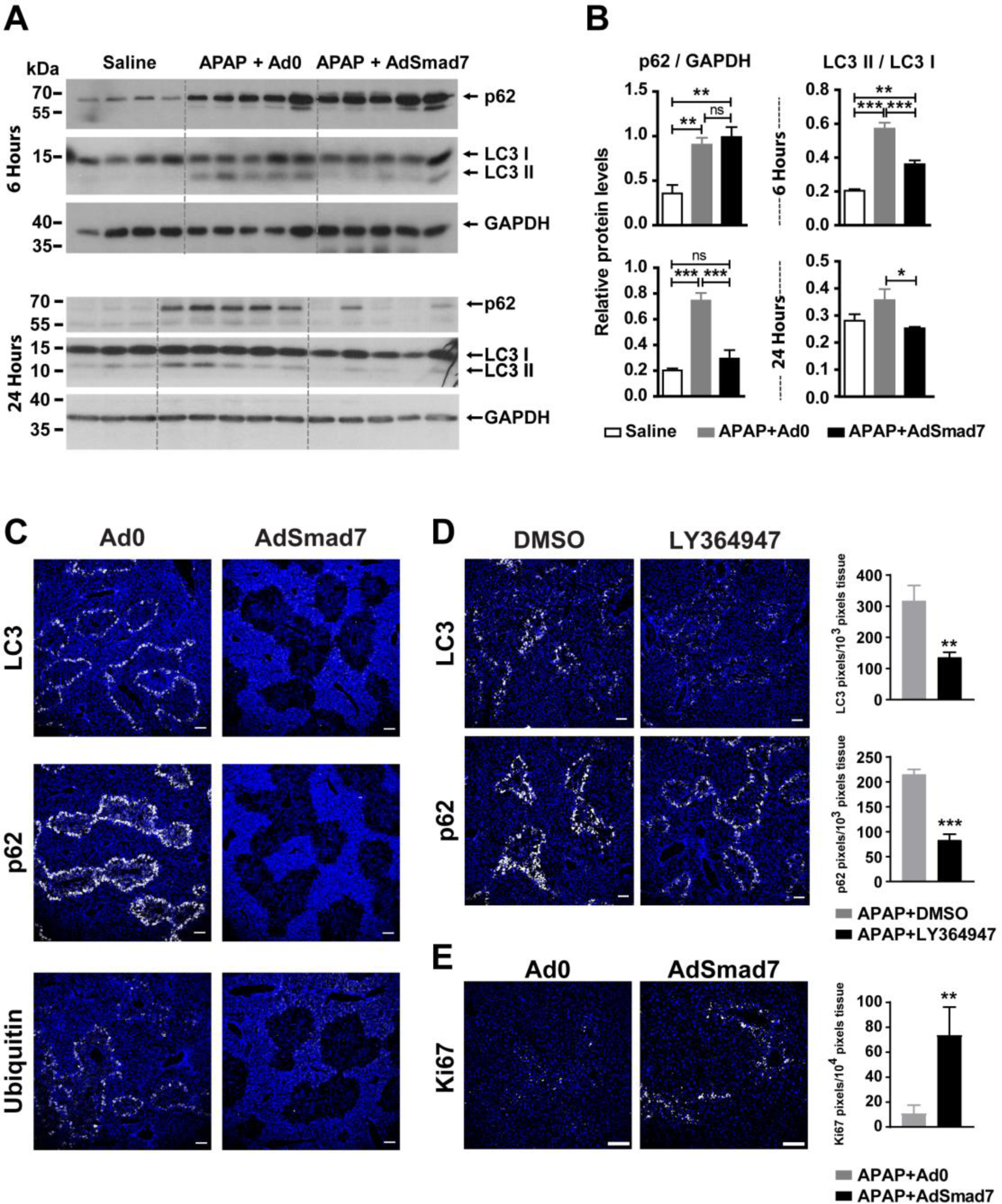
Inhibition of TGFβ-signaling blocks autophagy. (**A**) Western-blot analysis of p62 and LC3 levels. (**B**) Quantitative analysis of p62, relative to GAPDH, or a LC3II/ LC3I ratio, in saline or adenovirus plus APAP-treated male animals, 6 and 24 hours post-APAP treatment. Data represent mean±SEM of 4-5 independent samples per group and analyzed using one-way analysis of variance with Bonferroni’s post-hoc test. (**C-D**) Representative images of hepatic tissue sections derived at 24 hours from male animals treated with Ad0+APAP or AdSmad7+APAP (**C**), APAP plus DMSO or LY364749 (**D**), and stained for DAPI (blue) and LC3, p62 or Ubiquitin (white). LC3 and p62 fluorescence intensity in the perinecrotic areas was quantified using ImageJ software. Data are expressed as mean±SEM of 4-5 independent samples per group, analyzed using two-tailed unpaired Student’s t-test. (**E**) Representative images of Ad0- or AdSmad7-treated liver tissues 4 days post- APAP intoxication stained for Ki67 to visualize cycling hepatocytes. Ki67 expression was quantified by ImageJ-assisted analysis. Data are expressed as mean±SEM from 3- 4 animals per group analyzed using two-tailed unpaired Student’s t-test. Scale-bars in all images, 100μm.

**Figure 5.**
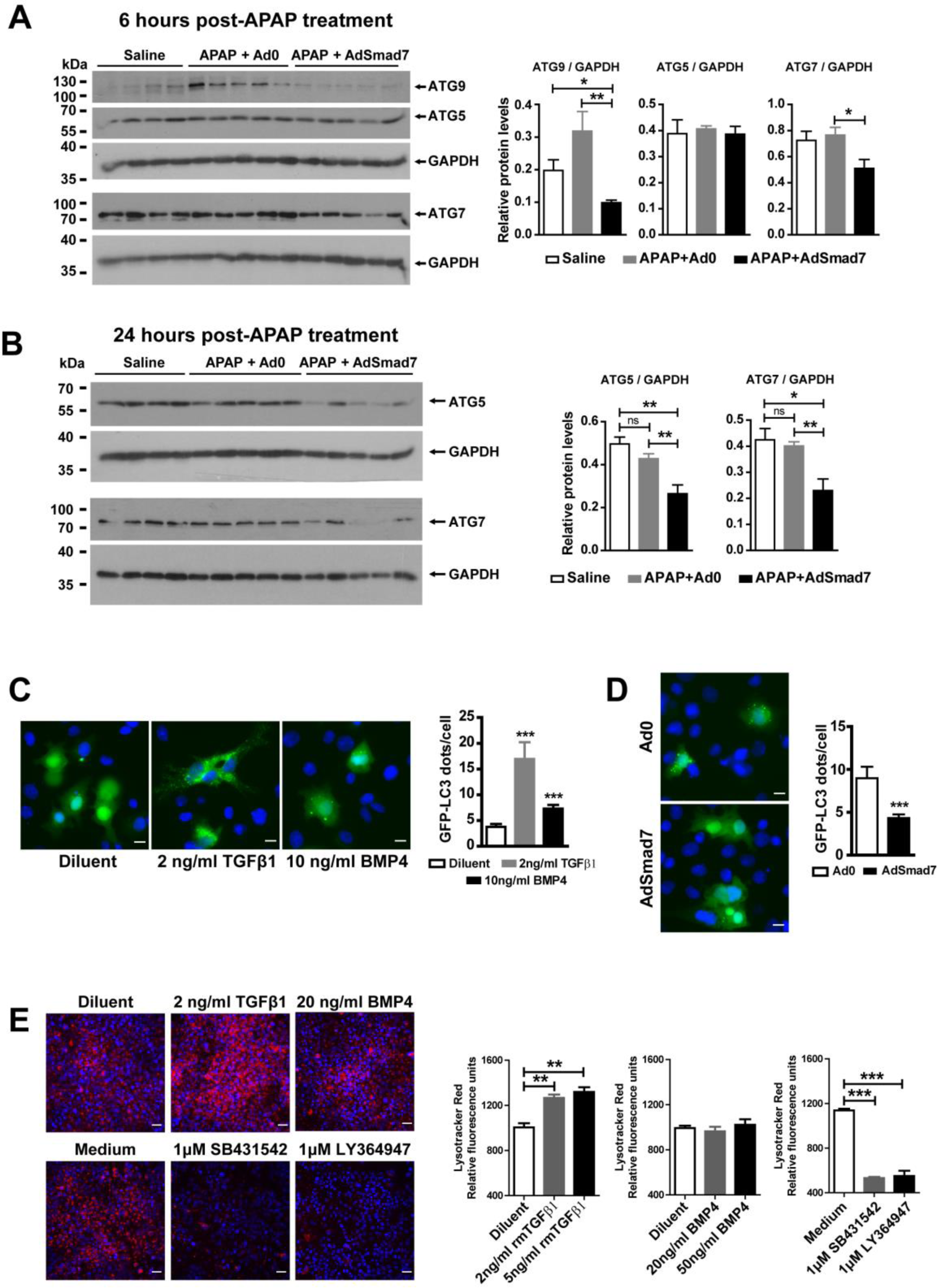
Alteration of TGFβ-signaling regulates autophagy and lysosomal activity. Western blot analysis of ATG5, ATG7, and ATG9 in saline or adenovirus plus APAP-treated male animals, 6h (**A**) and 24h (**B**) after APAP-administration. Data are expressed as mean±SEM of 4-5 independent samples per group and were analyzed using one-way analysis of variance with Bonferroni’s post-hoc test. (**C-D**) GFP-LC3+ punctuated autophagosomes detected by fluorescence microscopy, in Hepa1-6 cells treated with: diluent, 2ng/ml TGFβ1 or 10ng/ml BMP4 (**C**), Ad0 or AdSmad7 (**D**) for 24h (scale-bars, 20μm). Numbers of GFP-LC3+ puncta/cell were quantified in ∼400 cells/sample. Data are expressed as mean±SEM and analyzed using one-way analysis of variance with Bonferroni’s post-hoc test, or unpaired Student’s t-test for pairwise comparison. (**E**) Detection of acidic vacuoles by LysoTracker Red DNP-99 staining of Hepa 1-6 cells, treated for 24h with 2 or 5ng/ml TGFβ1, 20 or 50ng/ml BMP4, 1μΜ SB431542 or 1μΜ LY364947. Staining intensity was quantified using a fluorocount microplate reader and was normalized versus diluent or medium, respectively (scale-bars, 100μm). Data are expressed as mean±SEM of 3 independent samples/condition analyzed using one-way analysis of variance with Bonferroni’s post-hoc test

Immunostaining of tissue sections of APAP-intoxicated animals treated with AdSmad7 or LY364947 showed that the zones of intense autophagy at the borders of healthy and APAP-damaged tissues did not develop in the AdSmad7- or LY364947- treated animals (Figure 4C-D) illustrating the negative effect of TGFβ and/or BMP signaling inhibition on autophagy initiation.

To further substantiate the role of TGFβ and/or BMP signaling in hepatocyte autophagy, the mouse hepatoma Hepa1-6 cells were transfected with an LC3-eGFP plasmid and formation of cytoplasmic autophagic puncta was analyzed upon treatment with TGFβ1 and BMP4 ligands, or AdSmad7. Both ligands increased the number of LC3-eGFP**^+^** puncta, with TGFβ1 being the most potent inducer and AdSmad7 exhibiting an opposite effect (Figure 5C-D). Interestingly, only TGFβ1 could increase lysosomal activity which was inhibited by the addition of SB431542 and LY364947 (Figure 5E).

To better understand the accelerated healing of APAP-intoxicated, AdSmad7-treated livers, tissue sections were stained for Ki67 to visualize the proliferative status of hepatocytes. Substantially higher number of Ki67^+^ hepatocytes was detected in the AdSmad7-treated livers, explaining the accelerated tissue healing (Figure 4E).

The apparent discordance between severity of histopathology and speed of recovery in Smad7-treated animals further prompted us to investigate the death mechanism(s) involved. Twenty-four hours post-APAP administration, staining for HMGB1 (to monitor necrosis) and cleaved Caspase-3 (to monitor apoptosis) demonstrated absence of classical necrotic or apoptotic phenotype in AdSmad7-treated tissues (Figure 6A). Notably, EM analysis confirmed absence of autophagosomes in cells located at the borders of healthy and damaged tissue (Figure 6B). Instead, these cells displayed a paraptosis-like phenotype that involved enlarged cytoplasmic vacuoles, swollen mitochondria and absence of nuclear fragmentation or apoptotic-bodies. Moreover, CHOP, an endoplasmic reticulum stress-regulated protein and known inducer of paraptosis (33), was elevated both at 6 and 24h post-treatment (Figure 6C-D). It is reported that paraptosis is characterized by maintenance of cell membrane integrity (34). Accordingly, despite the aggravated histopathology of APAP+AdSmad7 treated animals (Figure 3E-F), at 6h post-APAP treatment, serum ALP, GGT, SGOT/AST and SGPT/ALT levels were significantly lower than those measured in APAP+Ad0 treated group and marginally higher than the saline-treated control group (Figure 6E). Moreover, APAP+Ad0 treated animals exhibited robust mobilization of inflammatory cells within damaged areas, activation of alpha smooth muscle actin (aSMA) positive stellate cells (Supplementary Figure 3A) and induction of CD45^+^ liver sinusoidal endothelial cells (Supplementary Figure 3B). Both latter cell-types are known players in liver regeneration (35, 36). Interestingly, Smad7 co-expression attenuated all the above reparatory responses. The attenuation of immune response in the AdSmad7- treated livers was further illustrated by increased hepatic mRNA levels for the anti-inflammatory cytokine *Il10* and decreased levels for the pro-inflammatory cytokines *Il13, Il4* and *Tnf* (Supplementary Figure 4). Collectively, the above data indicate that upon suppression of TGFβ signaling, APAP-intoxicated hepatocytes die passing through a stage with characteristics resembling paraptosis that is not accompanied by increased levels of hepatic enzymes in plasma.

**Figure 6.**
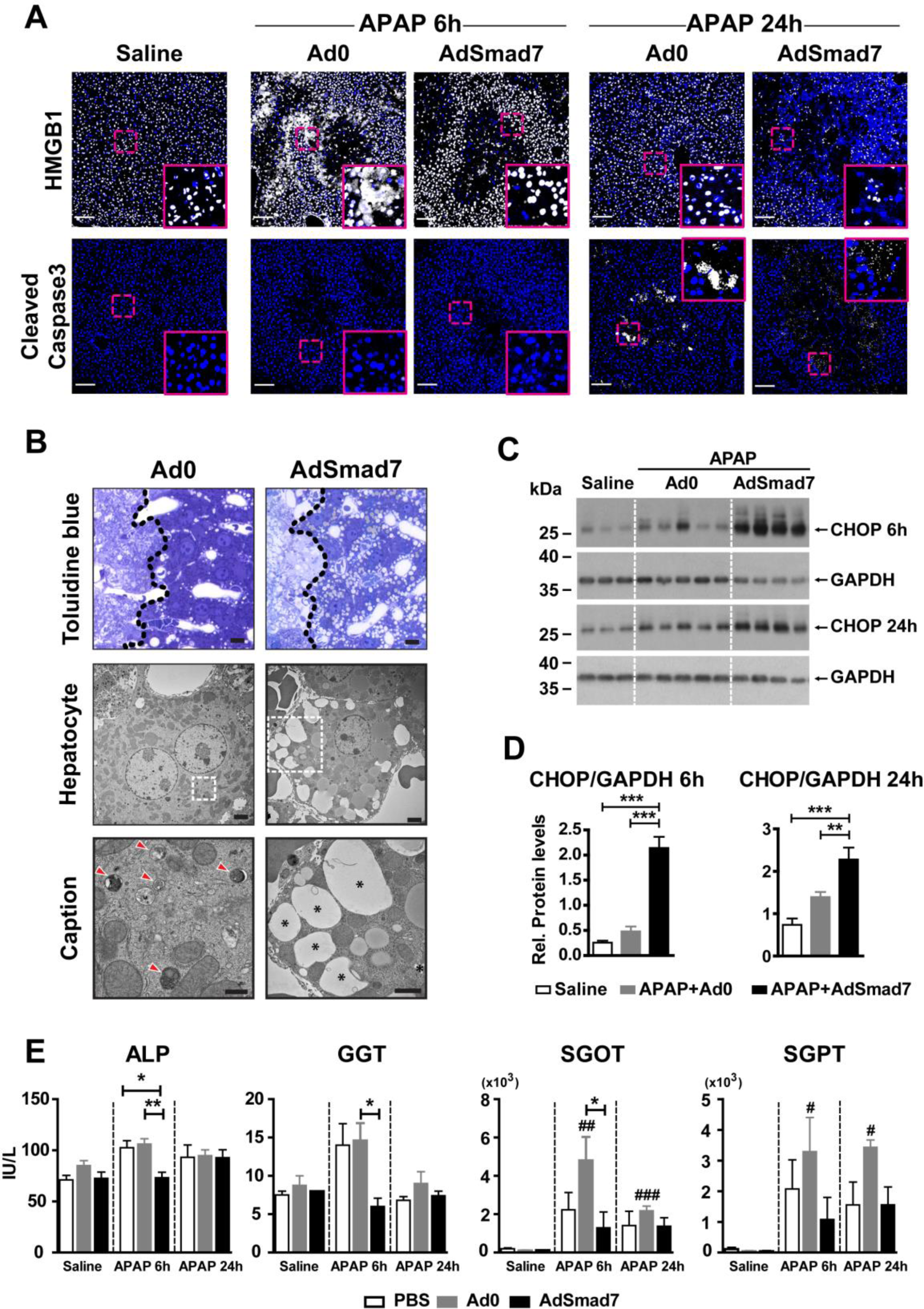
Smad7 over-expression in the liver of APAP-treated animals downregulates apoptotic markers and induces a paraptosis-like phenotype. (**A**) Representative images of hepatic tissue sections derived from PBS- or adenovirus-treated male animals, 6 and 24h following APAP or saline administration, stained for DAPI (blue), and HMGB1 or cleaved caspase-3 (white). Scale-bars, 100μm. Solid-line windows represent higher magnification of the areas shown with dashed-line squares. (**B**) Toluidine staining of liver sections, at the borders (dashed line) between healthy and damaged tissue (scale-bars, 20μm). Abundant autophagosomes (indicated by red arrowheads) or paraptosis-like vacuoles (indicated by asterisks) are detected by electron microscopy 24h following APAP administration, in the Ad0- or Smad7- treated animals, respectively (scale-bars, 2μm). Captions represent higher magnifications of areas marked by dashed-line windows in the middle column. Scale- bars, 1μm. (**C-D**) Representative Immunoblots for CHOP protein in control or adenovirus plus APAP-treated animals are shown in the left panels (Gapdh was used as loading control). Quantitative analysis of CHOP protein levels relative to Gapdh at 6 and 24h following APAP treatment is shown in the right panels. Data are expressed as mean±SEM of 4-5 independent samples/group, analyzed using one-way analysis of variance with Bonferroni’s post-hoc test. (**E**) Serum levels of ALP, GGT, SGOT, SGPT, in PBS- or adenovirus-treated male animals, 6 and 24h after APAP or saline administration. Data are expressed as mean±SEM of 5 independent samples/group, analyzed using one-way analysis of variance with Bonferroni’s post hoc-test.

### Transcriptional response of cultured primary hepatocytes following TGFβ1 and/or BMP4 stimulation highlights autophagy as a prominent biological process

Since reporter expression pointed out hepatocytes as prominent targets of TGFβ and/or BMP signaling, hepatospheres from primary mouse hepatocytes were cultured for 24h with recombinant TGFβ1, BMP4, both TGFβ1 and BMP4, or diluent, and their transcriptomes were analyzed through next-generation sequencing (RNA-seq). The transcriptome of livers treated with control or Smad7-expressing adenoviral vectors was also characterized, to enrich for putative direct Smad-targets and highlight possible mechanisms mediating the aggravating effect of Smad7.

BMP4 treatment led to significant upregulation of 2,780 and downregulation of 2,804 genes, whereas, upon TGFβ1 treatment, 1,488 genes were upregulated and 1,434 were downregulated. 27.5% of BMP4-responsive genes and 52.6% of TGFβ1-responsive genes were regulated similarly by both ligands (Figure 7A). A fraction of differentially expressed genes (∼5% for BMP4 and ∼10% for TGFβ1) were oppositely regulated when cells were stimulated with either ligand. However, a surprising interplay between the two ligands was revealed in cultures co-stimulated by TGFβ1 and BMP4. A striking characteristic of these cultures was the “dominance” of TGFβ1 over BMP4, as exemplified by the reversion of a substantial number of BMP4- regulated events back to baseline (1,669 upregulated and 1,677 downregulated genes). Comparison between ligand (*in-vitro*) and Smad7 (*in-vivo*) regulated transcriptomes demonstrated that Smad7 downregulated 345 genes that were upregulated by TGFβ1 or BMP4and upregulated 443 genes downregulated by both ligands (Figure 7B). We consider these gene sets as enriched in putative direct canonical Smad targets.

**Figure 7.**
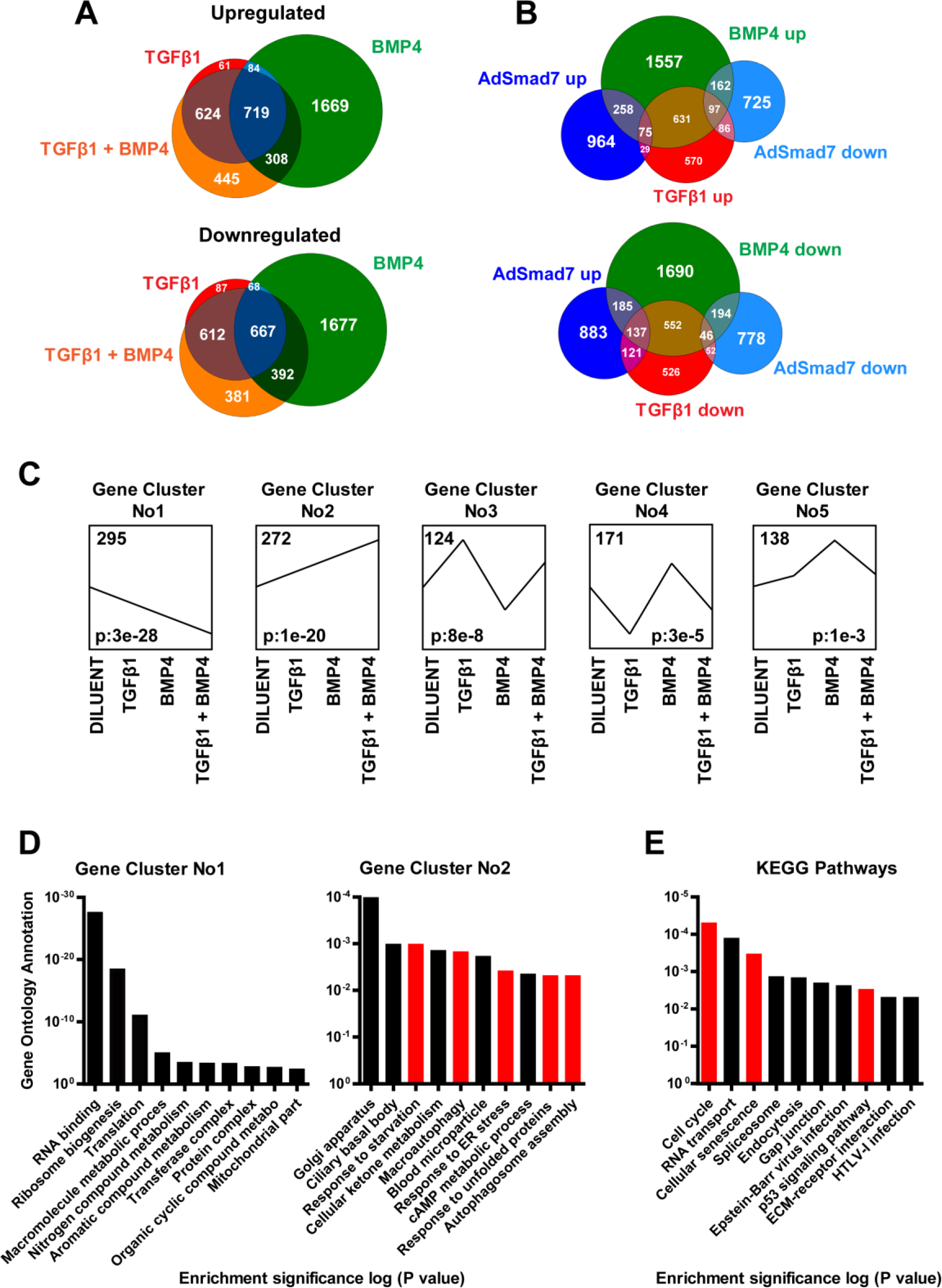
Transcriptomic analysis of TGFβ1 and/or BMP4 treated hepatospheres and AdSmad7-treated livers. (**A**) Venn-diagrams demonstrating overlapping patterns of upregulated or downregulated genes in hepatospheres treated for 24h with diluent, 5ng/ml rmTGFβ1, 20ng/ml rmBMP4, or a combination of both. Comparisons were done versus diluent-treated cultures. For each experimental group 3-4 independent samples derived from 8 hepatospheres each, were analyzed. (**B**) Venn- diagrams displaying the overlap between the RNA-Seq data of the AdSmad7- (n= 4) *vs* Ad0-treated (n= 4) livers (24h) or ligand- *vs* diluent-treated hepatospheres (up: upregulated, down: downregulated genes). For each ligand-treated experimental group, 3-4 independent samples derived from 8 hepatospheres each, were analyzed. (**C**) The five (5) most significant Gene Cluster Profiles of the ligand-stimulated hepatospheres identified by STEM analysis. Numbers at the upper left corner indicate the number of genes within the cluster and numbers at the lower-left corner the respective statistical significance (p-values). (**D**) The ten (10) most significant GO Annotations for Gene Clusters 1 and 2. Red bars highlight Annotations related to autophagy and ribosome biogenesis. (**E**) The ten (10) most significant KEGG pathways for the BMP4 uniquely downregulated genes. Red bars highlight Annotations related to cell cycle and senescence.

Gene ontology analysis of pathways activated by TGFβ1 or BMP4 (Table 1) demonstrated that some of them were shared between the ligands (for example, EIF2 Signaling*, Fatty acid β-oxidation, ketogenesis), some were regulated selectively by either ligand (for example “Oxidative phosphorylation”* and “Cholesterol Biosynthesis”* for TGFβ1, and “Regulation of Actin based Motility by Rho”* and “Bile Acid Biosynthesis” for BMP4) and some were oppositely regulated (for example “Integrin Signaling”, “Ethanol Degradation II”, “PPARα/RXRα Activation”). Several of these pathways were regulated by Smad7 oppositely to the ligands (indicated above by *). Interestingly, amongst the top pathways oppositely regulated by ligands and Smad7 were those related to autophagy (upregulated by ligands, downregulated by Smad7), translation-initiation, and ribosome-biogenesis (downregulated by ligands, upregulated by Smad7) (Figure 7C-D). Moreover, focused analysis of the genes uniquely downregulated by BMP4 (and normalized in the presence of TGFβ1) highlighted senescence as one of the top BMP4 downregulated pathways (Figure 7E, Supplementary Figure 5).

**Table 1:**
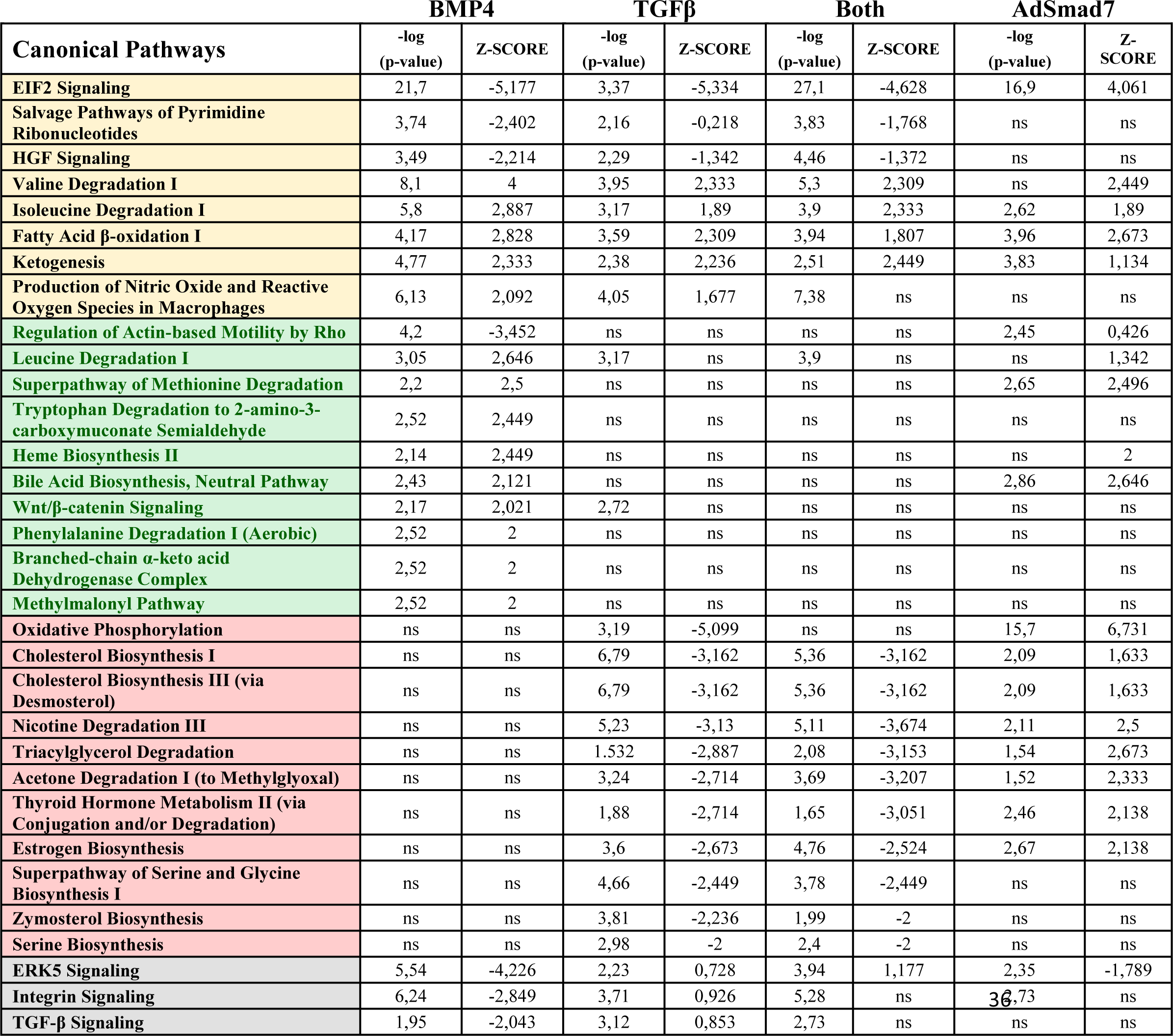
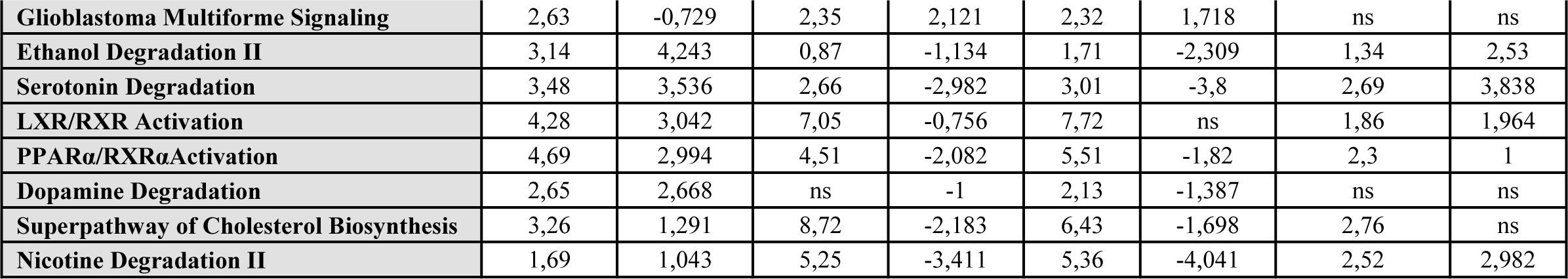
Bioinformatics analysis of pathways activated by TGFβ1/ BMP4 in hepatospheres, or AdSmad7-treated livers. Yellow color represents pathways commonly regulated by TGFβ1 and BMP4, green color represents pathways selectively regulated by BMP4, red color represents pathways selectively regulated by TGFβ1 and gray color represents pathways oppositely regulated by TGFβ1 and BMP4.

### The autophagy regulator *Trp53inp2* is upregulated by TGFβ-BMP signaling

Among all known regulators of autophagy found in the datasets (Figure 8A), *Trp53inp2*, a rate-limiting factor for initiation of autophagy (37), emerged as the top positively regulated gene by TGFβ1 and BMP4 and negatively regulated by Smad7 at the mRNA level. Treatment of Hepa1-6 cells revealed independently a similar and dose-dependent regulation of *Trp53inp2* mRNA levels by TGFβ1, BMP4, and Smad7 (Figure 8B). Accordingly, stronger nuclear TRP53INP2 immunostaining was detected in the pericentral, reporter positive, hepatocytes of saline-treated control livers (Figure 8C). TRP53INP2 immunostaining was substantially increased across the lobules in APAP-treated animals (Figure 9A-B, and Supplementary Figure 6A). Moreover, the punctuated cytoplasmic staining, a sign of TRP53INP2 activation and recruitment in the autophagic process, was detected selectively in the RFP^+^/eGFP^+^ zones of the APAP-treated animals, at the borders between damaged and intact tissues (Figure 8C). Importantly, concurrent Smad7 over-expression and APAP treatment abolished selectively the punctuated cytoplasmic TRP53INP2 distribution (Figure 9C). Interestingly, levels of ectopically expressed Smad7 declined faster than expected for adenovirus-mediated expression, dropping by almost 60% 24h post APAP treatment. This could be due to a selective loss of the AdSmad7-transduced hepatocytes and could account for the apparent absence of detectable reduction in nuclear TRP53INP2 immunostaining in the surviving hepatocytes at 24h post-APAP treatment. Thus, in addition to the transcriptional regulation of the *Trp53inp2* gene, it appears that TGFβ and/or BMP signaling could *in-vivo* regulate activation and/or cytoplasmic translocation of TRP53INP2 at the protein level.

**Figure 8.**
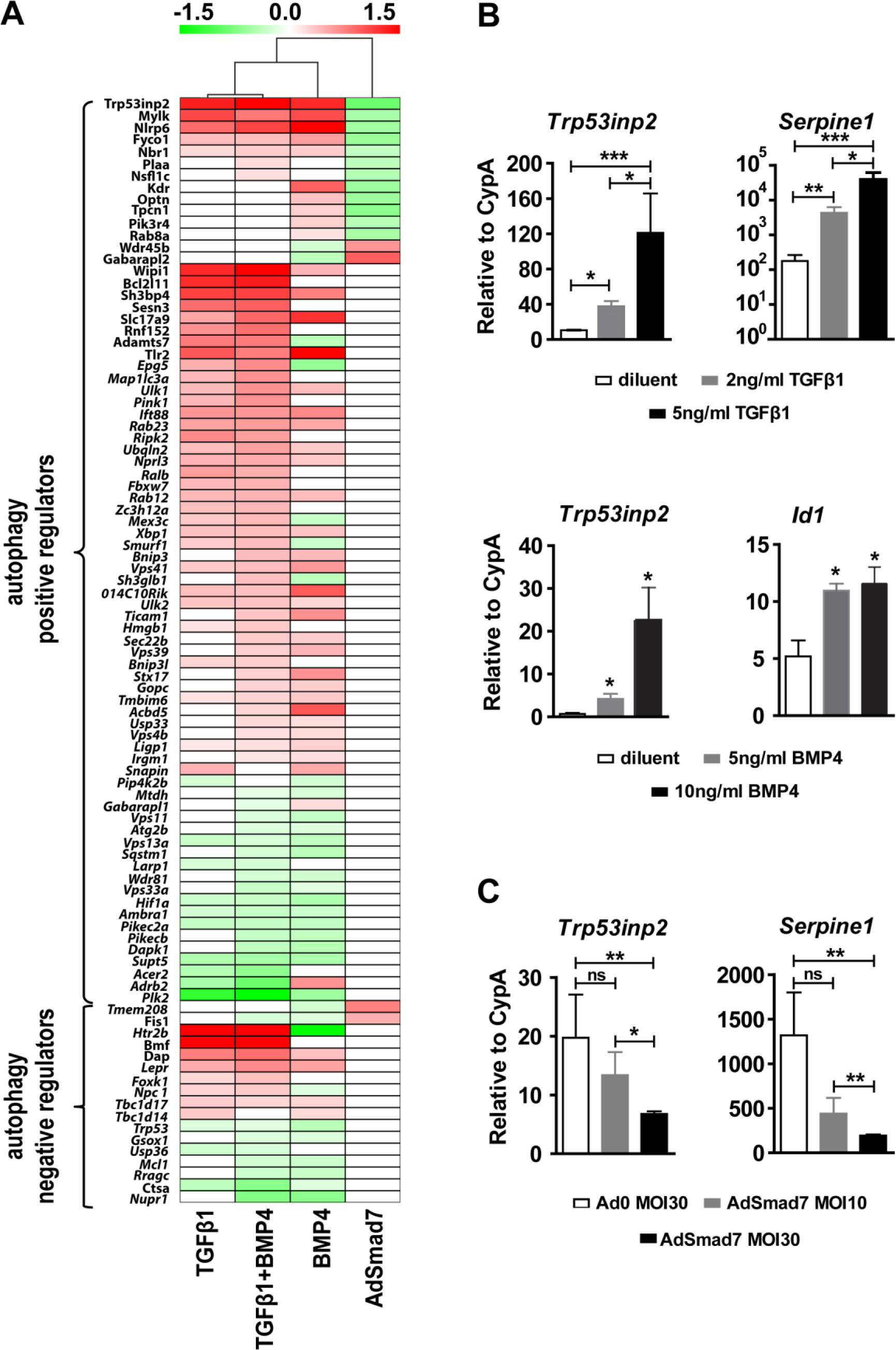
TGFβ/BMP pathways converge on the autophagy regulator *Trp53inp2*. **(A)** Heat-map of the log2 Fold Change values of autophagy related genes in the ligand-treated hepatospheres or AdSmad7-treated livers, based on the GO:0016238 (autophagy). **(B-C)** RT-qPCR analysis of *Trp53inp2, Serpine1* and *Id1* mRNA levels (relative to Cyclophilin A levels), 24h after treatment of Hepa1-6 cells with TGFβ1 or BMP4 (**B**), Ad0 or AdSmad7 (**C**). Data are expressed as mean±SEM of three independent samples per group analyzed using one-way analysis of variance with Bonferroni’s post-hoc test.

**Figure 9.**
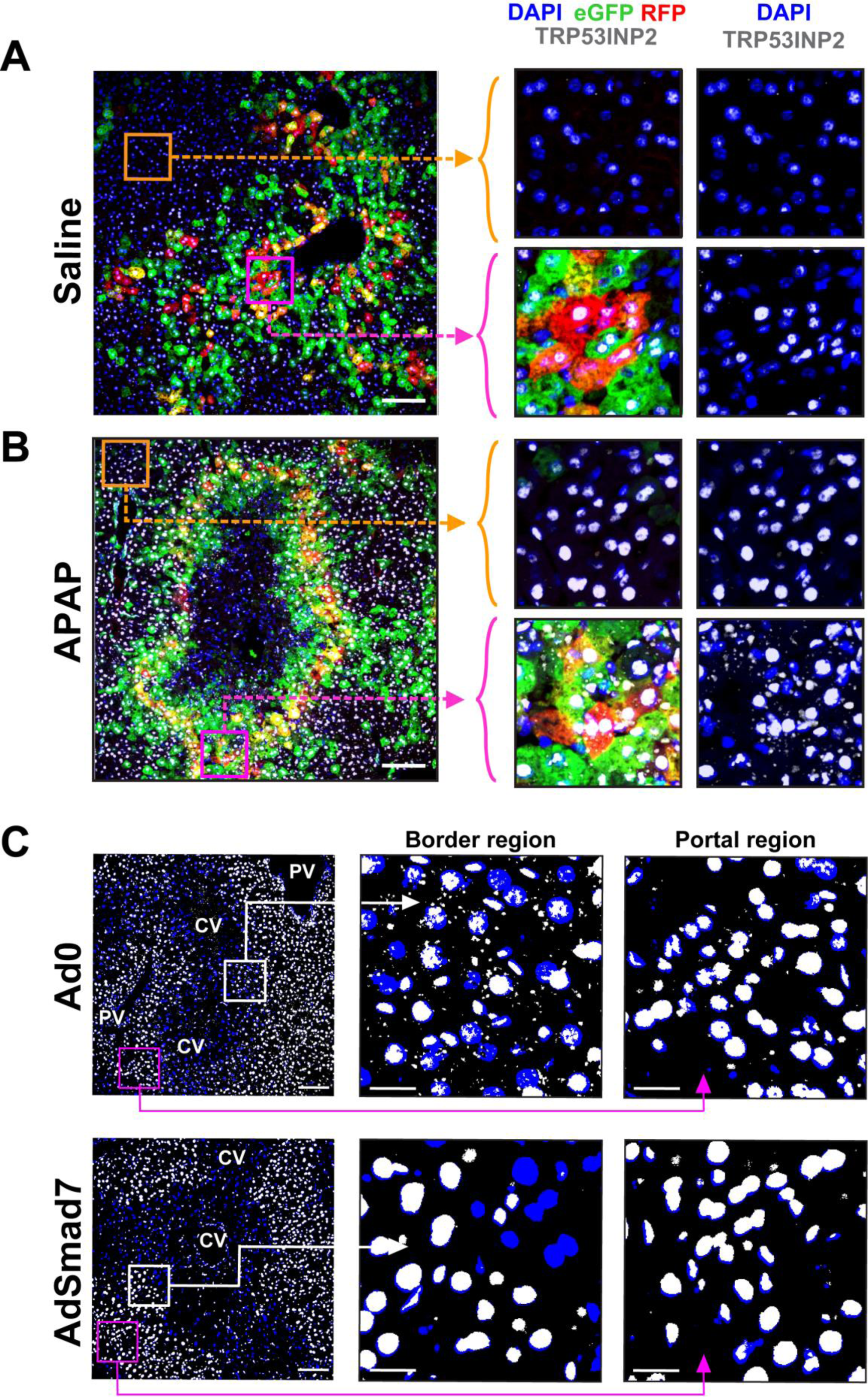
Association between TGFβ/BMP pathways, autophagy, and Trp53inp2. (**A-B**) Representative confocal images of hepatic tissue sections from saline (**A**) or APAP-treated female animals (**B**) 24h post-treatment (scale-bars, 100μm), stained for eGFP (green), RFP (red), TRP53INP2 (white), and DAPI (blue). The right panels show magnification of selected areas from the reporter positive (magenta line window) or reporter negative (orange line window) hepatic zones. (**C**) Representative confocal images of liver tissue sections from adenovirus-treated male animals, 24h post-APAP administration, stained for TRP53INP2 (white), and DAPI (blue). White and magenta windows represent tissue areas at the borders of the injured tissues or the portal region proximal, undamaged tissues, respectively (scale-bars, 100μm). Central Vein (CV), Portal Vein (PV).

Given the rapid increase of both TGFβ and BMP reporter activity during early postnatal period and the well-established role of autophagy in remodeling neonatal hepatic tissues (38), we investigated whether TRP53INP2 is regulated in a similar manner during this stage. Indeed, the kinetics of TRP53INP2 expression during hepatic early postnatal development and in particular its cytoplasmic punctuated pattern (Supplementary Figure 6B) mirrored the kinetics of reporter expression and Smad phosphorylation (Figure 1B-D), strongly suggesting that TGFβ-superfamily signaling could regulate autophagic process during neonatal liver development as well.

## Discussion

Despite substantial progress in the characterization of multiple components and processes in the TGFβ-superfamily signaling system, the way by which cells within a tissue integrate and decode the complex TGFβ-superfamily generated signaling input to evoke the desired physiological responses, in particular the way they perceive the interplay between TGFβ/Activin- and BMP-signaling branches, still remains unclear. The overarching objective of this study was to investigate the functional interplay between TGFβ/Activin and BMP signaling in the context of hepatic physiology and pathophysiology. To this end, a novel double-reporter animal model was developed to visualize *in-situ* activation of canonical TGFβ/Activin- and/or BMP-signaling. This unique tool enabled us to observe at the single-cell level, the dynamic, highly coordinated, stage- and context-specific, spatiotemporal patterns of TGFβ/Activin and/or BMP reporter expression during development and adult liver pathology and demonstrate the functional interconnectivity of the two TGFβ-superfamily signaling branches in health and disease. Reporter expression was detected in the liver already as early as embryonic day E14.5, the earliest time point analyzed, increased gradually during the prenatal period and exhibited explosive activation during the early postnatal stage, when the liver undergoes dramatic adaptation to self-feeding mode (39). Development towards adulthood was associated with gradual and coordinated contraction of the reporter zones towards the central veins of the lobules. Establishment of tight Glutamine Synthetase peri-venus zones before the complete contraction of the reporter zones in central regions may suggest that TGFβ- superfamily does not play a role in establishing liver zonation but rather is under the influence of mechanisms that control it, such as, for example, the Wnt pathway (40). Activation of the TGFβ superfamily system and pattern of the reporter expression were also analyzed in the context of acute pathology using the APAP overdose mouse model. Increased pSMAD2 and pSMAD1/5/8 levels detected in protein extracts from APAP-intoxicated livers demonstrated robust activation of both signaling branches. Correspondingly, immunostaining of hepatic tissues from APAP-treated reporter animals demonstrated strong activation of both reporters in zones demarcating the borders between healthy and damaged tissues, 24h post-drug administration. The hepatocytes in these TGFβ and BMP reporter double-positive zones were characterized by enhanced immunostaining for autophagic markers and were found by EM to be rich in autophagosomes. Anatomically similar zones characterized also by intense macroautophagy using EM have been reported in a prior study of APAP- treated mouse livers (41). The accumulation of molecules involved in autophagy, such as p62, LC3II and ubiquitin at the borders of the injured tissues can be interpreted in two different ways. They could indicate either a blockade of autophagic flux or, alternatively, an overwhelmed activation of the autophagic machinery (31, 42). Previous findings argue in favor of the latter alternative. The capacity of APAP to activate protective autophagic responses in hepatocytes is well documented (31, 32). Moreover, increased levels of LC3II or GFP-LC3-positive puncta have been previously described in APAP-treated mouse hepatocytes *in-vitro* and *in-vivo* (31, 43). Therefore, the most probable explanation for the observed accumulation of the autophagy markers and glutathionylated protein adducts is saturated autophagic machinery. This phenomenon is more evident at the outer borders of the damaged areas, most likely because the hepatocytes within these areas are severely damaged and beyond repair.

In the current study, we demonstrate that at 6h post-APAP treatment, blockade of TGFβ signaling by Smad7, leads to decreased LC3II levels, while the protein levels of p62 remain high. Moreover, reduced protein levels of the autophagy regulators ATG7 and ATG9 detected at this time point, probably account for the inhibition of LC3 lipidation and the formation of LC3II^+^ early autophagosomes (44) and the blocking of autophagy machinery at early stages of autophagosome biogenesis (45). On the contrary, levels of ATG5, a protein involved at a stage preceding LC3 lipidation and autophagosome closure, are unaltered.

The link between TGFβ/BMP signaling and autophagic activity was further supported by analyzing the transcriptomes of primary hepatocytes stimulated *in-vitro* by TGFβ1 and/or BMP4. Bioinformatics analysis highlighted autophagy amongst the top regulated pathways. While autophagy has been previously associated with either APAP intoxication or TGFβ signaling (46–48), this is the first report where coordinated and cooperative activation of both BMP and TGFβ pathways is implicated in the context of APAP-induced autophagy in the liver. A mechanistic explanation for this cooperation was provided by the analysis of known autophagy-related molecules, which highlighted TRP53INP2, a rate-limiting factor for autophagosome assembly (49), as a putative mediator of the TGFβ/BMP effects on the autophagic activity of APAP-intoxicated hepatocytes. TRP53INP2 immunostaining is found higher in reporter positive hepatocytes located in the vicinity of the central veins and is substantially increased upon APAP intoxication. Moreover, *Trp53inp2* mRNA levels and most importantly, the cytoplasmic distribution of TRP53INP2 appear to be positively regulated by TGFβ/BMP signaling and suppressed by Smad7.

Interestingly, TGFβ-superfamily signaling associated changes in TRP53INP2 expression and subcellular localization can be detected also during early postnatal development. This period which is characterized by robust TGFβ/BMP reporter activation is also known to entail intense remodeling of the hepatic parenchyma driven to a great extent by autophagy. These findings raise the fascinating possibility that TGFβ/BMP cooperative regulation of autophagy process via TRP53INP2 could represent a fundamental role of this signaling pathway in the context of both liver physiology and pathophysiology.

Inhibition of the TGFβ signaling system either by Smad7 or pharmacological inhibitors, besides demonstrating its crucial role for APAP-induced autophagy alterations, led to additional and quite intriguing findings. Specifically, overexpression of Smad7 in the APAP-treated livers or systemic administration of selective TGFβ/Activin inhibitors revealed dual and at first glance contradicting roles for the TGFβ-superfamily signaling system. Hence, its inhibition was associated initially with exacerbation of APAP-induced liver histopathology, consistent with a protective role, which was followed by a surprising acceleration of tissue recovery at later time-points, consistent apparently with a detrimental role for the TGFβ- superfamily system. Similar “dual” behavior has been previously demonstrated in p53 deficient animals (50), where deficiency in this key cell-cycle “gatekeeper”, aggravated APAP-induced acute hepatic injury but enhanced the subsequent regenerative capacity of the liver.

Accelerated tissue recovery of APAP-intoxicated liver upon inhibition of TGFβ receptor signaling has been recently described by Bird *et al*. (24). This study provided evidence for TGFβ-signaling activation in hepatocytes located at the borders of areas of APAP-induced necrosis, and implicated it to the induction of a senescence-associated secretory-phenotype that could spread pathology to the entire organ causing total liver failure. Using selective TGFβ receptor inhibitors, the authors demonstrated a reduction in senescence markers and acceleration of liver regeneration that protected animals from APAP overdose. Interestingly, despite the agreement on the accelerated healing associated with inhibition of TGFβ-signaling Bird *et al.,* did not observe the enhanced histopathology induced by TGFβ inhibitors in our study. Reasonably, the final outcome of APAP-induced liver injury must be determined by the balance between the severity of tissue damage and the effectiveness of tissue- reparatory and tissue-regenerative mechanisms. It is possible that the relative contribution of the tissue-reparatory or regenerative mechanisms recalled under given circumstances to handle liver injury could vary in a context-dependent manner. For example, in younger individuals, where the regenerative machinery is still very efficient, inhibition of TGFβ signaling could further shift the balance in favor of regeneration and, overall, might be beneficial despite the induction of a feebler tissue repair response. On the contrary, in the elderly, where the regenerative capacity has declined and cannot be enhanced effectively, sacrificing the TGFβ-driven reparatory machinery could have detrimental consequences. Indeed, age has been highlighted as an important factor in APAP-induced liver injury, with young people being relatively less sensitive and a more potent regenerative capacity being one of the suggested explanations (51). Whether the discrepancy between our study and Bird’s *et al.* (24), is due to differences in the age of animals analyzed (6-8 weeks, *versus* ∼12 weeks) warrants further investigation especially given the postulated therapeutic utility of TGFβ inhibitors for APAP intoxication.

Another interesting observation in our study involves the form of programmed cell-death hepatocytes undergo upon APAP administration in the presence or absence of canonical TGFβ-superfamily signaling and the involvement of a paraptosis-like death as the prominent form of cell loss in AdSmad7-treated animals. Paraptosis is characterized by exuberant cytoplasmic vacuolation driven by a stressed endoplasmic-reticulum. Interestingly, the increased vacuolation is also evident in the histology of APAP+TGFβ receptor inhibitor treated tissues in the study of Bird *et al*. (24). Moreover, *Atg7* deficient hepatocytes form similar endoplasmic reticulum derived enlarged cytoplasmic vacuoles (52), further supporting a possible link between paraptosis-like phenotype and suppression of autophagy.

TGFβ and BMP signaling pathways behave in a cooperative manner as far as autophagy is concerned; however, the interplay between these signaling branches appears to be more complex. The prevailing notion regarding their interaction is that they are antagonistic, however, examples of redundant or cooperative behavior have also been reported (53, 54). In line with these observations, pathway analysis in the current study unveiled novel functions regulated by TGFβ1 and/or BMP4 in an antagonistic, cooperative, or selective manner. The antagonistic component of the TGFβ/BMP interplay became evident in hepatocytes treated simultaneously with both ligands. TGFβ1 substantially dominated the transcriptional profile of the hepatospheres, reverting many of the BMP4-regulated genes to baseline levels. Interestingly, similar observations have been made in *Xenopus* embryos, where TGFβ-induced Connective Tissue Growth Factor (CTGF) could physically interact with BMP4 and neutralize it (55). Therefore, expression of CTGF could provide a mechanistic explanation for the TGFβ1 over BMP4 dominance. Indeed, in the hepatocyte cultures used herein, CTGF was induced by TGFβ (log2FoldChange: 2.02), suppressed by BMP4 (log2FoldChange: -1.07) and importantly was also upregulated in the presence of both ligands (log2FoldChange: 2.16).

A small number of processes appear to be oppositely regulated in hepatocytes by TGFβ or BMP stimulation *in-vitro*. Interestingly, whereas TGFβ receptor inhibition can suppress APAP-induced hepatocyte senescence (24), as shown herein, stimulation of hepatocytes with BMP4 only, can downregulate transcription of several senescence-associated genes. Thus, BMP signaling could function as an inhibitor of senescence in the eGFP positive regions of the hepatic lobules, restraining thus the secretory senescence phenotype that emanates from the double-reporter positive zone that demarcates the damaged tissues. The overall balance between TGFβ- and/or BMP-driven protective or damaging responses in healthy or APAP-injured tissues is summarized in Figure 10. It is worth noting that despite the strong similarities between human and mouse liver/hepatocytes regarding their response to APAP intoxication, some differences have been described by *in-vitro* and patient-based studies (56, 57). Therefore, confirmation of some of the conclusions of the current study with human cells/tissues is warranted.

**Fig. 10.**
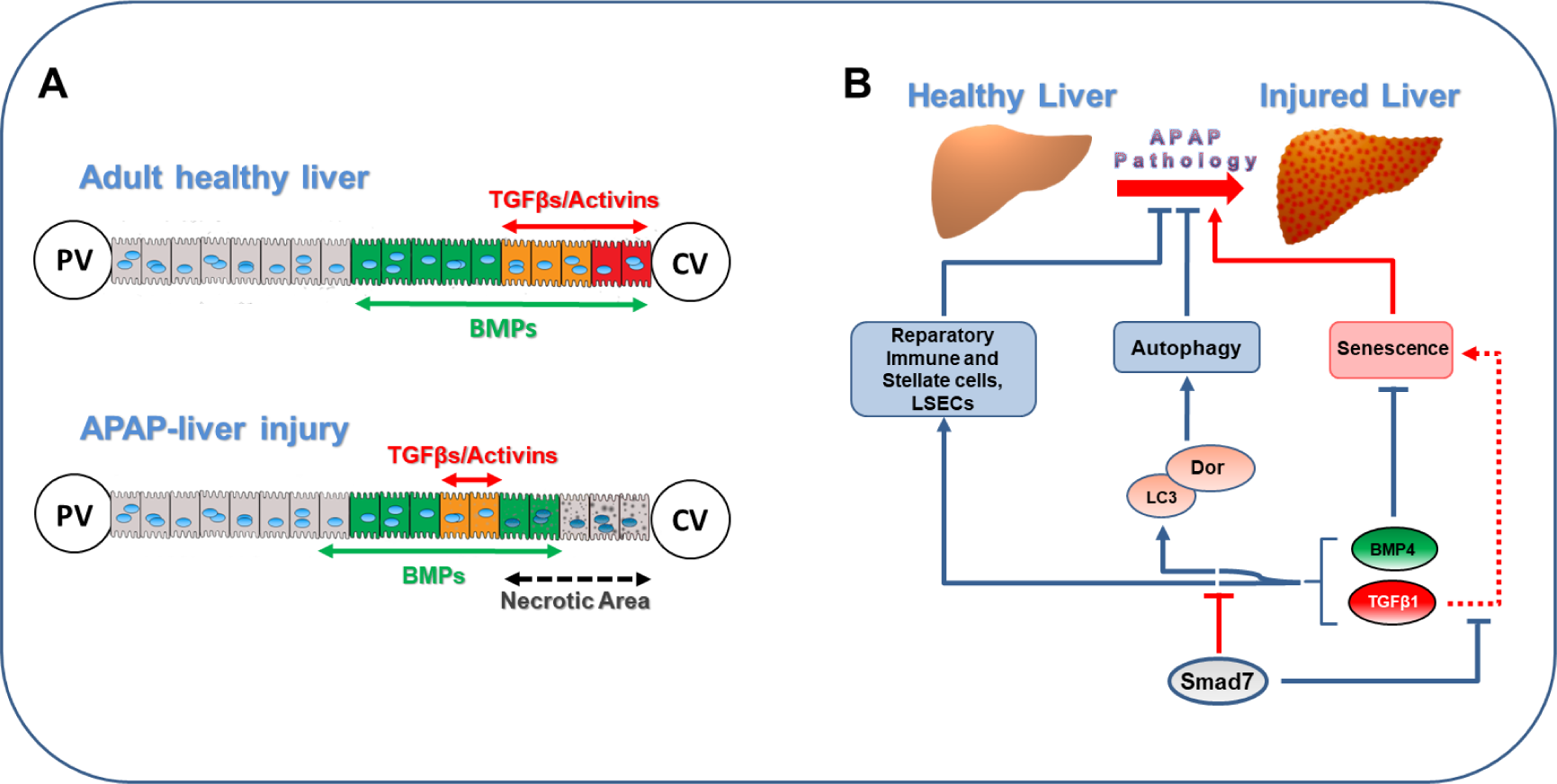
Schematic representation summarizing the TGFβ superfamily regulated responses in healthy or injured liver, upon APAP intoxication. **(A)** Tissue distribution of the TGFβ- and BMP-responding cells in healthy and injured liver. **(B)** Model depicting the putative dual TGFβ/BMP-signaling switch that regulates mobilization of reparatory cell populations, autophagy and senescence. Blue connection lines highlight beneficial effects; red lines represent harmful effects and dotted line information from the literature (Bird *et al. 2018*).

A novel and crucial aspect of the current study from both basic and translational points of view is the illustration of the strong interconnectivity of the TGF- superfamily signaling system. This is exemplified by the dynamic and coordinated activation of the two reporters in spatiotemporally overlapping zones in health and disease and the dynamic balance of the transcriptomes induced by TGFβ1 and/or BMP4 stimulation. The identification of gene sets that are regulated in a redundant, selective, or opposite way by either ligand in solitude and the dominance of TGFβ1 over BMP4 during co-stimulation with both ligands should be taken into consideration while interpreting the consequences of genetic or pharmacological inactivation/inhibition of individual receptors, ligands or regulatory components which are specific to either signaling branch. Henceforth, the findings and tools described herein could guide the study of the TGFβ superfamily signaling in liver pathophysiology and aid the development of novel and/or safer therapeutic interventions for acute and chronic liver diseases.

## Materials and Methods

### Preparation of reporter construct and generation of transgenic lines

A TGFβ-responsive (TRE), RFP expressing reporter construct, similar to the previously described BMP-responsive construct CMVe-BRE-eGF (30) was produced utilizing the TGFβ/Activin reporter vector (CAGA)_12_-MLP-Luc (58) by inserting the CMVe element upstream of the (CAGA)_12_-MLP element, and by introducing at both ends of the construct the chicken β-globin HS4 insulator to protect the reporter cassette from silencing effects. The HS4 insulator was isolated from the pVZc-HS4 vector (gift from Dr. Georgios Vasilopoulos, BRFAA). The Luciferase sequence was replaced by the mRFP1 (monomeric Red Fluorescent Protein1) sequence that was isolated from the pCx-mRFP plasmid (28). The final construct, hereafter referred to as “TRE-RFP” was sequenced, released from the plasmid backbone, cleaned and injected in the pronucleus of C57BL/6 fertilized oocytes, giving rise to six TRE-RFP founders (Mouse Transgenics Facility, Erasmus MC, Rotterdam, The Netherlands). Mice carrying the transgene were identified by Southern blot and PCR, using the primers 5’-GCCTCCTCCGAGGACGTCATCAAG-3’ (sense) and 5’- CGCCGGTGGAGTGGCG GCCC-3’ (antisense) (59) to amplify the 610 bp mRFP1 sequence. The line with the highest transgene expression (F6) was selected for further analysis.

### Animals

TGFβ/Activin and BMP double reporter animals (TRE-BRE mice) were established by crossing the BMP-responsive BRE-eGFP mouse line (30) with the TGFβ- responsive TRE-RFP line F6. The animals were maintained as heterozygotes, in C57BL/6 background. Reporter and C57BL/6 control animals were housed at the Animal House Facility of the Biomedical Research Foundation of the Academy of Athens (Athens, Greece), in individually ventilated cages under specific pathogen free conditions, on a 12:12 hours light:dark cycle and *ad-libitum* access to food and water. All procedures for care and treatment of animals were in full compliance with FELASA and ARRIVE recommendations and were approved by the Institutional Committee on Ethics of Animal Experiments and the Greek Ministry of Agriculture.

### Liver perfusion and isolation of mouse primary hepatocytes

Primary hepatocytes were isolated from two months old females. To minimize diurnal variation, isolation was initiated at 9-10 a.m. The abdomen of anaesthetized animals was opened to expose the portal vein, followed by opening of the chest and cannulation of the central vein with a 20G plastic cannula (Hospira, G717-A01), through the *vena cava*. Perfusion was initiated with 40ml of pre-warmed (37°C) Liver Perfusion Medium (Gibco, 17701) supplemented with 1% Penicillin-Streptomycin (Gibco, 15070-063) at a flow rate of 1 ml/min. The portal vein was cut to allow flow of the perfusate. Liver was perfused for 20-25min to completely remove blood. Perfusion continued with 50ml pre-warmed (37°C) Liver Digest Medium (Gibco, 17703) supplemented with extra 250μl of 100mg/ml collagenase (Sigma-Aldrich, C5138), at a flow rate of 1ml/min. Thereafter, the liver was transferred into a 10cm Petri dish, the bladder was removed, 10-15ml of Hepatocyte Wash Medium (Gibco, 17704) supplemented with Penicillin-Streptomycin was added and the tissue was minced with forceps. Homogenized tissue was filtered through a 100μm cell strainer and primary hepatocytes were collected after a series of three centrifugations in 20ml Hepatocyte Wash Medium, at 50g, for 5min, at room temperature. Cell viability was estimated by trypan blue staining and was generally greater than 85%.

### Cell cultures and Hepatosphere formation

Hepa1-6 cells (ATCC® CRL-1830™) were cultured in DMEM (Gibco, 41966-052) supplemented with 10% FBS (Gibco, 10500) and 1% Penicillin/Streptomycin. Six hours before the addition of recombinant mouse TGFβ1 Protein (R&D Systems, 7666-MB-005), recombinant mouse BMP4 Protein (R&D Systems, 5020-BP-010) [diluted in 0.1% BSA, 4mM HCL, dH2O-diluent], SB431542 (Sigma-Aldrich, S4317) or LY364947 (Tocris Bioscience, 2718) [diluted in DMSO (Sigma-Aldrich, D2650)-vehicle], the culture medium was replaced with fresh medium containing 2% FBS. Cells were transfected with Polyethylenimine-PEI Transfection Reagent (Polysciences, 23966-2) carrying the LC3-eGFP plasmid (60) (a kind gift from Dr. Tamotsu Yoshimori, Osaka University, Japan) and infected with Ad0 or AdSmad7 at a Multiplicity Of Infection (M.O.I)=30. Cells were collected 24h later for mRNA analysis or fluorescence quantitation. Alternatively, cells were incubated for 20min with 75nM LysoTracker Red (Molecular Probes, L-7528), and fixed with 4% paraformaldehyde (Merck, 104004), prior to fluorescent quantitation.

Hepatospheres were formed by culturing purified mouse hepatocytes at a density of 5x10^5^ cells/ml in 60 μl hanging drops in Williams E medium (Gibco, A12176), supplemented with 10% FBS (Gibco, 10500), 100 nM Dexamethasone (Sigma-Aldrich, D4902), 1% Penicillin/Streptomycin, 0.5 mg/ml Gentamycin (Biochrom, A2712) and 1% Insulin-Transferrin-Selenium (Gibco, 41400-045). Where indicated, hepatospheres were cultured in the presence of recombinant mouse TGFβ1 or BMP4, and/or the selective TGFβ receptor inhibitors SB431542 and LY364947. Untreated and diluent or vehicle-treated hepatospheres were used as controls. Cells were cultured at 37°C in a humidified atmosphere containing 5% CO2.

### Histology

Organs and tissues were surgically removed, rinsed in PBS and placed in PBS containing 4% paraformaldehyde for 24h at 4°C. Subsequently, the tissues were placed in PBS with 30% sucrose for 24h at 4°C, rinsed with PBS, embedded in OCT (Shandon Cryomatrix) and kept at -80°C until use. Cryostat sections 10µm thick (Leica CM1950) were placed on poly-L-lysine slides (Menzel Glaser, 19352000), and stored at -80°C. Antigen retrieval with 10mM Sodium Citrate buffer, pH: 6.0 for 30min in 80°C water-bath followed by 30min incubation on ice was performed before immune-staining. After blocking with the appropriate donkey or goat normal serum, tissues were incubated over-night at 4°C with the following primary antibodies: rat anti-RFP (Antibodies-Online, ABIN334653), chicken anti-GFP (Abcam, ab13970), rabbit anti-Glutamine Synthetase (Antibodies-Online, ABIN398821), rabbit anti-Cleaved Caspase-3 (Cell Signaling, 9661), rabbit anti-p62 (MBL, PM045), rabbit anti-LC3 (MBL, PM036), rabbit anti-Ubiquitin (Dako, Z0458), rabbit anti-HMGB1 (Abcam, EPR3507), rat anti-mCD45 (BD Biosciences, 550539), rabbit anti-alpha smooth muscle Actin (aSMA) (Abcam, ab5694), rabbit anti-Ki67 (Abcam, ab15580) and rabbit anti-TRP52INP2 (ThermoFisher, PA5-38729). Sections were washed and incubated for 60min with secondary antibodies: goat anti-rat Alexa594 (Molecular Probes, A11007), donkey anti-chicken FITC (Jackson Immuno-research, 703-096- 155) and donkey anti-rabbit Alexa 647 (Jackson Immuno-research, 711-606-152). The slides were mounted using the ProLong Gold Antifade Reagent with DAPI (Molecular Probes, P36931). Images were captured with a Leica DMRA2 fluorescent microscope equipped with Leica DFC320 and DFC350 FX digital cameras and a Leica TCS-SP5 confocal microscope (Leica Microsystems, Wetzlar, Germany). Images were processed and analyzed using Adobe Photoshop CS3 and ImageJ 1.41.

### Electron Microscopy, Toluidine Blue and Haematoxylin/Eosin staining

For transmission electron microscopy, mice were anesthetized and abdominal cavity was opened. The portal vein was connected to the perfusion system and animals were fixed for 5min with 2.5% glutaraldehyde in 0.1 M phosphate buffer, pH 7.4. The liver was removed in a petri dish containing fixative and dissected into 1-mm cubes. After subsequent buffer washes, samples were post-fixed in 1.0% osmium tetroxide for 1h on ice. After dehydration through graded ethanol series, each sample was infiltrated gradually in a mixture of Epon/Araldite resins diluted in propylene oxide and then embedded in fresh epoxy resin mixture. Finally, the specimens were allowed to polymerize at 60°C for 24h. To verify section quality and select areas of interest, 1- μm-thick sections were cut and stained with Toluidine Blue (1% with 2% borate in distilled water). Ultrathin sections were cut with a Diatome diamond knife at a thickness of 65 nm on a Leica EM UC7 ultramicrotome (Leica Microsystems, Vienna, Austria), were then mounted onto 300 mesh copper grids and stained with uranyl acetate and lead citrate. Sections were examined on a Philips 420 transmission EM at an acceleration voltage of 60 kV fitted with a Megaview G2 CCD camera (Olympus SIS, Münster, Germany). For Haematoxylin and Eosin (H&E) staining, livers were excised en bloc, submersed in 10% buffered formalin overnight, and processed for paraffin embedding and sectioning. Five (5) μm histological sections cut in the coronal plane, were loaded onto poly-L-lysine slides, deparaffinized in xylene and rehydrated in graded ethanol dilutions and distilled H_2_O and thereafter, stained with H&E. Images were captured with a Leica DMRA2 microscope equipped with Leica DFC320 and DFC350FX digital cameras. For morphometric analysis, H&E stained liver sections were color deconvoluted and a 40X final magnification picture/sample was recorded by two indented observers and used to compute mean group values for each of the stated measurements using the ImageJ 1.41 program.

### Quantification of Fluorescence signal

Local fluorescence maxima of RFP expression in hepatospheres and LC3-eGFP puncta in Hepa1-6 cells were determined using the “find maxima” function of the ImageJ software package. The sum of the signal intensities of the selected points was used for the quantitation of the overall RFP expression and the actual number of points (elements) per cell for the estimation of eGFP positive puncta. LysoTracker Red was quantified in fixed Hepa1-6 cells with the Fluorocount Microplate Reader (Packard, AF10000). Analysis of immunostaining for Ki67, TRP53INP2, p62 and LC3 in control and APAP-treated animals was performed with the ImageJ software package using the “adjust threshold” function. Large vessels and necrotic areas were excluded from the analysis. For p62 and LC3 only the peri-necrotic zones, where accumulation of these molecules occurs, were analyzed after manual selection of these zones with ImageJ.

### mRNA expression analysis by Real-Time PCR and RNA-Seq

For Real-Time quantitative PCR (RT-qPCR) analysis, total RNA was isolated using the TriReagent protocol (Sigma-Aldrich). cDNA synthesis was performed as previously described(61). RT-qPCR was performed using primer pairs and conditions selected with the “Primer-BLAST” software (62) as follows: for RFP, 5’- ATCCCCGACTACTTGAAG-3’ (sense) and 5’-CATGTAGGTGGTCTTGAC-3’ (antisense); eGFP, 5’-CATCTTCTTCAAGGACGAC-3’ (sense) and 5’- TTGTGGCTGTTGTAGTTG-3’ (antisense); *Id1*, 5’-GGCGAGATCAGTGCCTTG- 3’ (sense) and 5’-AAGGGCTGGAGTCCATCTG -3’ (antisense); *Serpine1* primers, 5’-CTCCGAGAATCCCACACAG-3’ (sense) and 5’- ACTTTGAATCCCATAGCATC-3’ (antisense); *Trp53inp2* (Dor), 5’- GGTATGCGGCTCCAGTTGT-3’ (sense) and 5’- CTATGGCAGTTTAAAGCCCTGG-3’ (antisense); *Gapdh* 5’- CCAGTATGACTCCACTCACG-3’ (sense) and 5’- CTCCTGGAAGATGGTGATGG-3’ (antisense); and Cyclophilin A, 5’- ACCGTGTTCTTCGACATCACG-3’ (sense) and 5’- CTGGCACATGAATCCTGGAATA-3’ (antisense). Results were collected with a Chromo4 RT-qPCR detector and analyzed with the Opticon Monitor3 software (Bio-Rad Laboratories). Relative levels of mRNA expression were normalized to *Gapdh* or Cyclophilin A and calculated according to the ΔΔCT method (63). For transcriptomic analysis, total RNA was extracted using the RNeasy® Micro kit (Qiagen, 74004) from hepatospheres, or using TriReagent (Sigma-Aldrich, T9424) from the Ad0 or AdSmad7 adenovirus-treated livers. cDNA libraries were produced using the TruSeq RNA kit (Illumina) and sequenced in the Greek Genome Center at BRFAA. Differential (pairwise) analysis of count data was estimated by the *DESeq2* method (64). Genes with BaseMean counts per library >10 and False Discovery Rate (FDR) <0.05 were selected for downstream analysis. Data were analyzed using QIAGEN’s Ingenuity® Pathway Analysis (IPA®, QIAGEN Redwood City), and the Short Time-series Expression Miner (STEM) software (65). Venn diagrams were designed using the Venn Diagram Plotter (Pacific Northwest National Laboratory, U.S. Department of Energy) or the R platform Euler package.

### Adenovirus production and administration and *invivo* LY364947 administration

Recombinant adenoviruses containing FLAG-tagged Smads were kindly provided by Dr. Kohei Miyazono (University of Tokyo, Japan) and Dr. Aristidis Moustakas (Uppsala University, Sweden). Control empty adenovirus (Ad0) was generated as previously described (66). Adenovirus propagation and titration were performed as previously described (67). A total of 5x10^9^ (Ad0 or AdSmad7) or 1x10^10^ (Ad0, AdSmad1 or AdSmad3) infectious particles diluted in 200μl PBS were injected intravenously in the mouse tail and livers were collected at the indicated time points. 10mg/kg of LY364947 or equal volume of DMSO was diluted in saline and administered intraperitoneally in a volume of 10μl/g mouse weight.

### Acetaminophen preparation and administration

Acetaminophen (Sigma-Aldrich, A7085) was diluted in warm saline and administered intraperitoneally in a volume of 10 μl/g mouse weight. Experiments were performed either with female mice (TRE-BRE reporter and C57BL6 wild-type), treated with 400 mg/kg APAP, or male C57Bl6 wild-type mice, treated with 250 mg/kg APAP, 16h after adenoviral intravenous administration or 1.5h after intraperitoneal LY364947 administration.

### Biochemical analysis in serum

Blood was collected by retro-orbital bleed and allowed to clot, followed centrifugation in a refrigerated centrifuge -4°C for serum separation and collection. Serum levels of Alkaline phosphatase (ALP), Gamma-Glutamyl Transferase (GGT), Glutamic-Oxaloacetic Transaminase (SGOT/AST) and Glutamic-Pyruvic Transaminase (SGPT/ALT) were measured by colorimetric assays (Medicon Hellas) using a Medilyzer biochemical analyzer (Medicon Hellas).

### Western Blot Analysis

For protein extraction, cells or organs were homogenized as previously described (61). Homogenized samples were centrifuged at 50.000g for 60min at 4°C and supernatants were stored at -80°C until use. Protein concentration was calculated with Bradford assay (68). Samples containing 30-40μg total protein were separated on 10- 13% SDS-polyacrylamide gels under reducing conditions and proteins were transferred onto nitrocellulose membranes. Total Protein bands that transferred in nitrocellulose membranes were rapidly detected by Ponceau S staining (0.2% Ponceau S, 1% Acetic acid), and staining removed by washing multiple times with 0.05% Tween20 in PBS (PBS-T). Membranes blocked with 5% non-fat milk in PBS-T or Tris-Buffered Saline TBS (TBS-T) (RT, 60min) and immunoblotted using the following primary antibodies (4°C, overnight): rabbit anti-pSmad1/5/8 (Merck Millipore, AB3848), rabbit anti-pSmad2 (Cell Signaling, 3101S), mouse anti-GAPDH (Chemicon, MAB374), mouse anti-β-actin (Sigma-Aldrich, A2228), rabbit anti-ATG9 (Calbiochem, ABC23), rabbit anti-ATG5 (Sigma-Aldrich, A0856), rabbit anti-ATG7 (Abgent, AP1813D), rabbit anti-p62 (MBL, PM045), rabbit anti-LC3 (MBL, PM036), rabbit anti-Ubiquitin (Dako, Z0458) and rabbit anti-CHOP (Santa Cruz, sc-575). For the detection of glutathionylated protein conjugates, a mouse anti-Glutathione antibody (Abcam, ab19534) was used under non-reducing conditions. Primary antibody binding was detected with goat anti-rabbit HRP (Jackson Immunoresearch, 111-035-003) or rabbit anti-mouse HRP (Jackson Immunoresearch, PAB0096) antibodies (RT, 90min). Bands were visualized by chemiluminescence using ECL^TM^ Western blotting Detection Reagent (Amersham Biosciences). Band intensity was measured by using the Gel Analyzer 1.0 software.

### Data availability

Gene expression data are available from Sequence Read Archive (https://www.ncbi.nlm.nih.gov/sra), accession number SUB6360391.

### Statistical analysis

Data were analyzed using one-way analysis of variance followed by Bonferroni’s post-hoc analysis or, two-tailed unpaired Student’s t-test, using GraphPad Prism. One asterisk (*) corresponds to statistical significance of P< 0.05, two asterisks (**) to P < 0.01, and three asterisks (***) to P < 0.001.

## Acknowledgments

Authors would like to thank Prof. C.L. Mummery (Leiden University, The Netherlands) for providing BRE-eGFP reporter animals, Dr. Peter Ten Dijke (Leiden University, The Netherlands), Dr. Georgios Vasilopoulos (BRFAA, Greece) and Dr. Anna-Katerina Hadjantonakis (Memorial Sloan Kettering Cancer Center, USA), providing plasmids critical for the construction of the TRE-RFP transgene, Prof. Frank Grosveld, John Kong-A-San and Dr. Alex Maas (Erasmus MC, The Netherlands) for TRE-RFP transgenic microinjections/generation, Dr. Tamotsu Yoshimori (Osaka University, Japan) for providing the LC3-eGFP plasmid and Drs E. Rigana and S. Pagkakis (Imaging Facility, BRFAA, Greece) for their support in confocal microscopy. Recombinant adenoviruses containing FLAG-tagged Smads were kindly provided by Dr. Kohei Miyazono (University of Tokyo, Japan) and Dr. Aristidis Moustakas (Uppsala University, Sweden). Dr. A. Moustakas is also thanked for critical reading of the manuscript.

## Abbreviations

APAP: Acetaminophen
aSMA: alpha Smooth Muscle Actin
ALP: Alkaline phosphatase
ATG5: Autophagy Related 5
ATG7: Autophagy Related 7
ATG9: Autophagy Related 9
BMP: Bone Morphogenetic Protein
BRE: BMP Responsive Element
CHOP: DNA Damage Inducible Transcript 3
CTGF: Connective Tissue Growth Factor
CypA: cyclophilin A
DMSO: Dimethylsulfoxide
eGFP: enhanced Green Fluorescent Protein
EM: Electron Microscopy
FCS: Fetal Calf Serum
GAPDH: Glyceraldehyde-3-Phosphate Dehydrogenase
GGT: Gamma-Glutamyl Transferase
GS: Glutamine Synthase
HMGB1: High Mobility Group Box 1
LC3: Microtubule Associated Protein 1 Light Chain 3 Alpha
PPARα: Peroxisome Proliferator Activated Receptor α
RFP: Red Fluorescent Protein
RXRα: Retinoid X Receptor α
SGPT/ALT: Serum Glutamic Pyruvic Transaminase/Alanine Transaminase
SGOT/AST: Serum Glutamic-Oxaloacetic Transaminase/Aspartate Aminotransferase
SQSTM1/P62: Sequestosome 1 or p62
TGFβ: Transforming Growth Factor beta
TNFα: Tumor Necrosis Factor α
TRE: TGFβ Responsive Element
TRP53INP2: Tumor Protein P53 Inducible Nuclear Protein 2

