## Supplemental Material for "Coordinated activation of both TGFβ and BMP canonical pathways regulates autophagy and tissue regeneration in acetaminophen induced liver injury"

**Supplementary Figure 1**

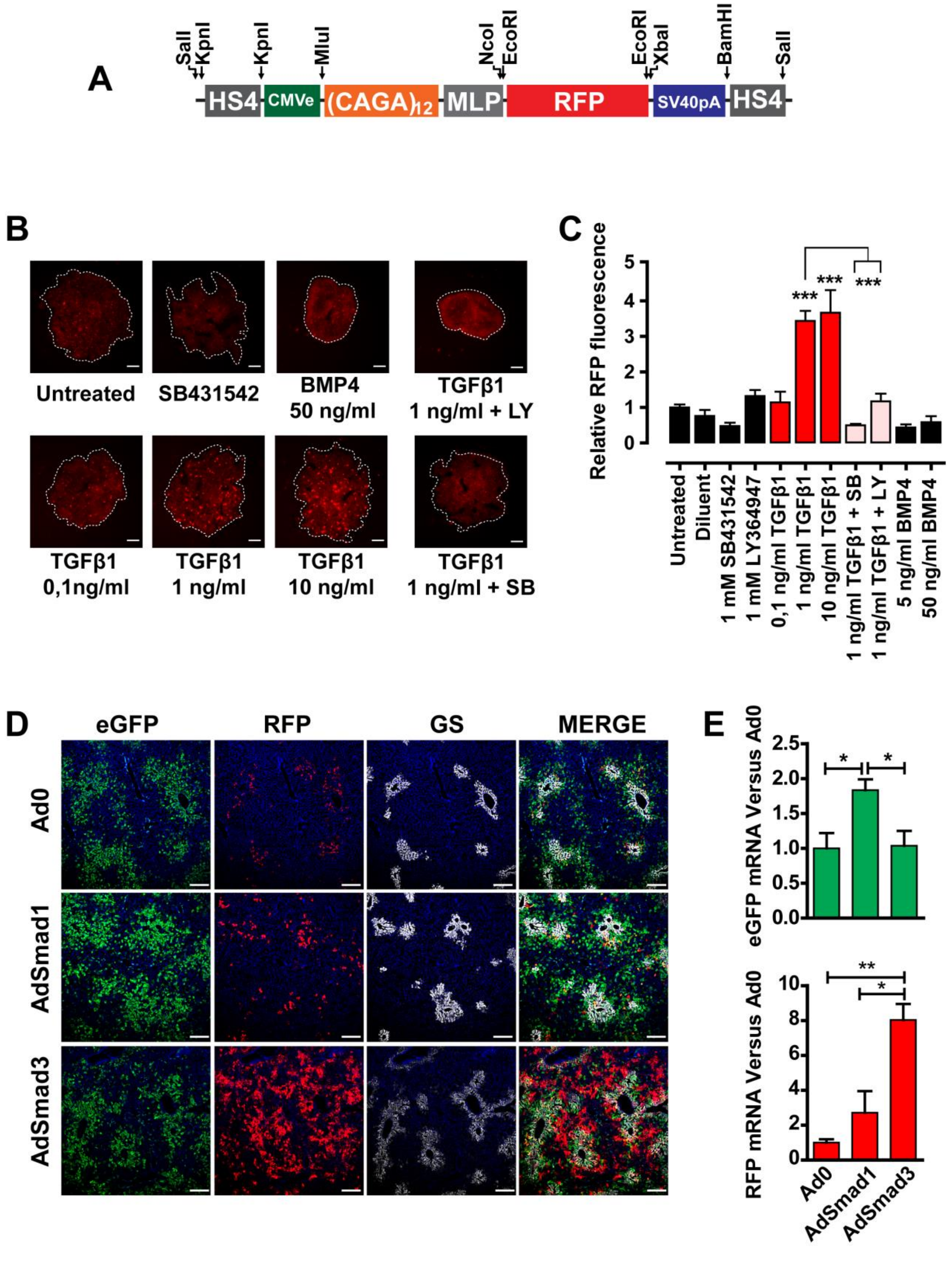

### Supplementary Figure 2

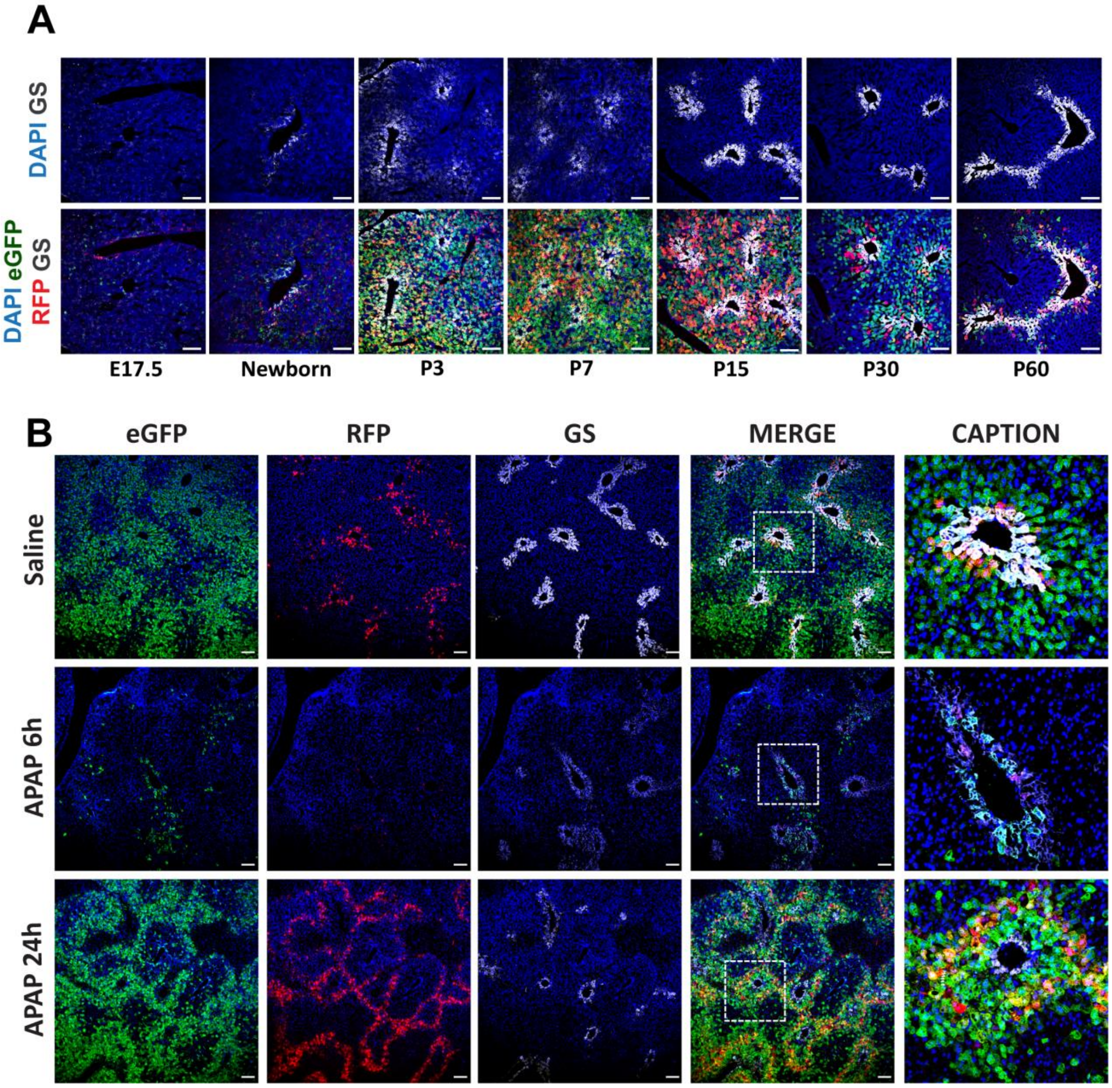

### Supplementary Figure 3

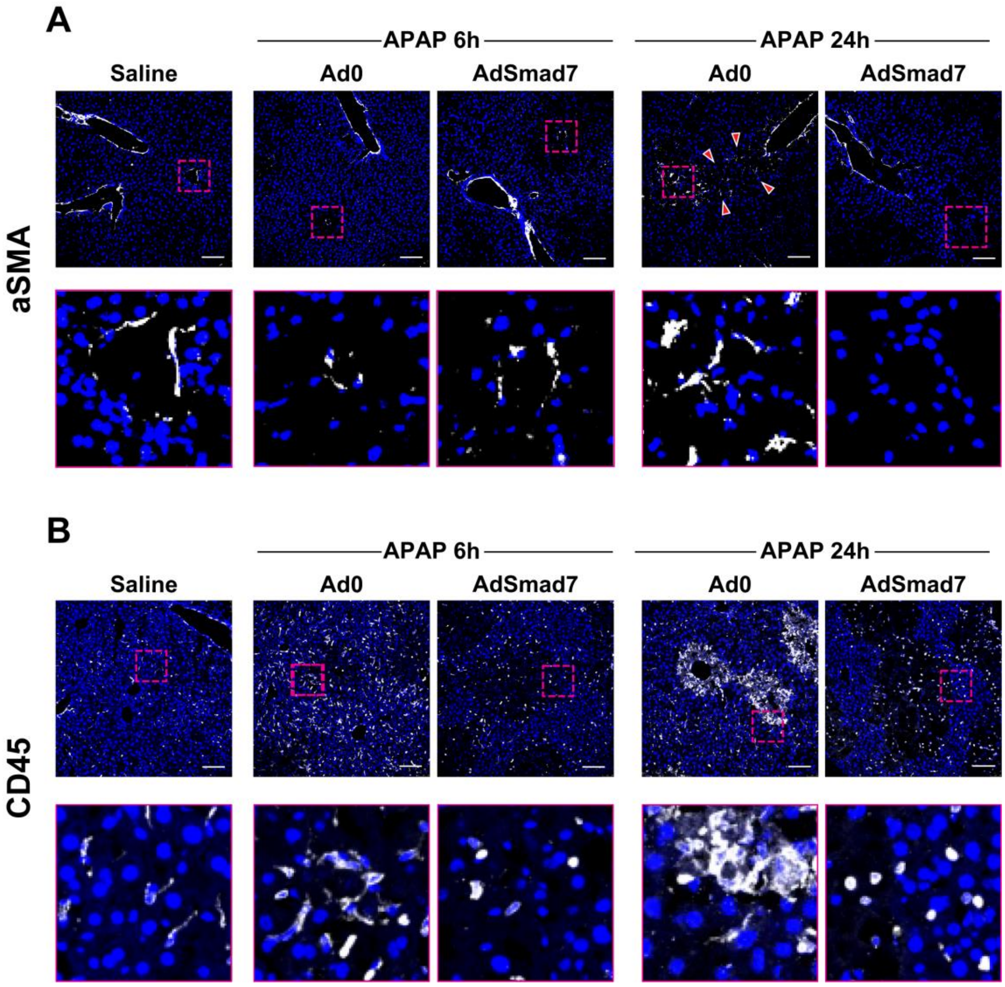


### Supplementary Figure 4

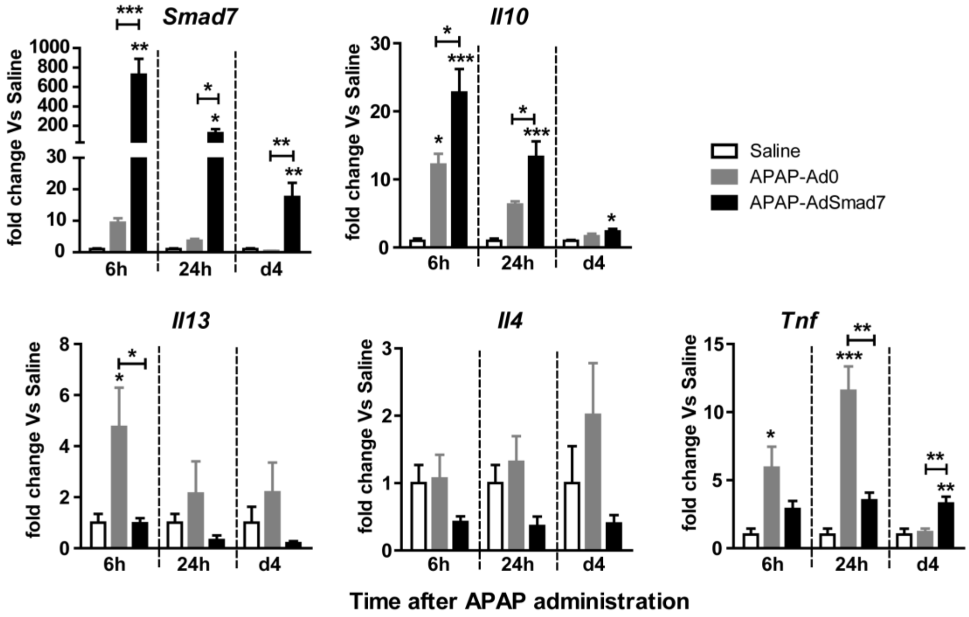

**Supplementary Figure 5**

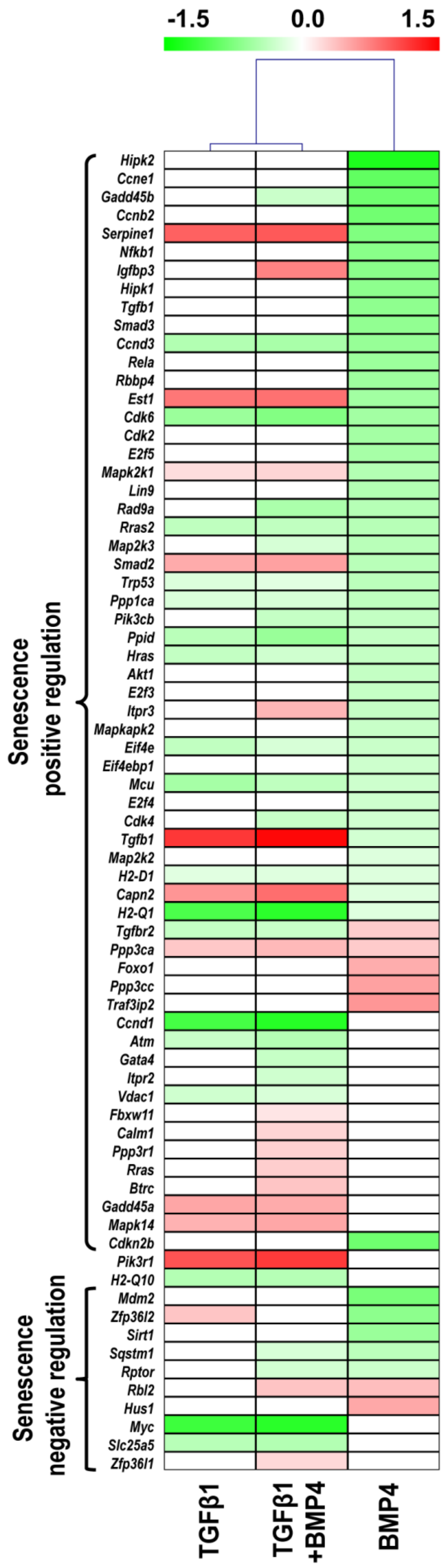

**Supplementary Figure 6**

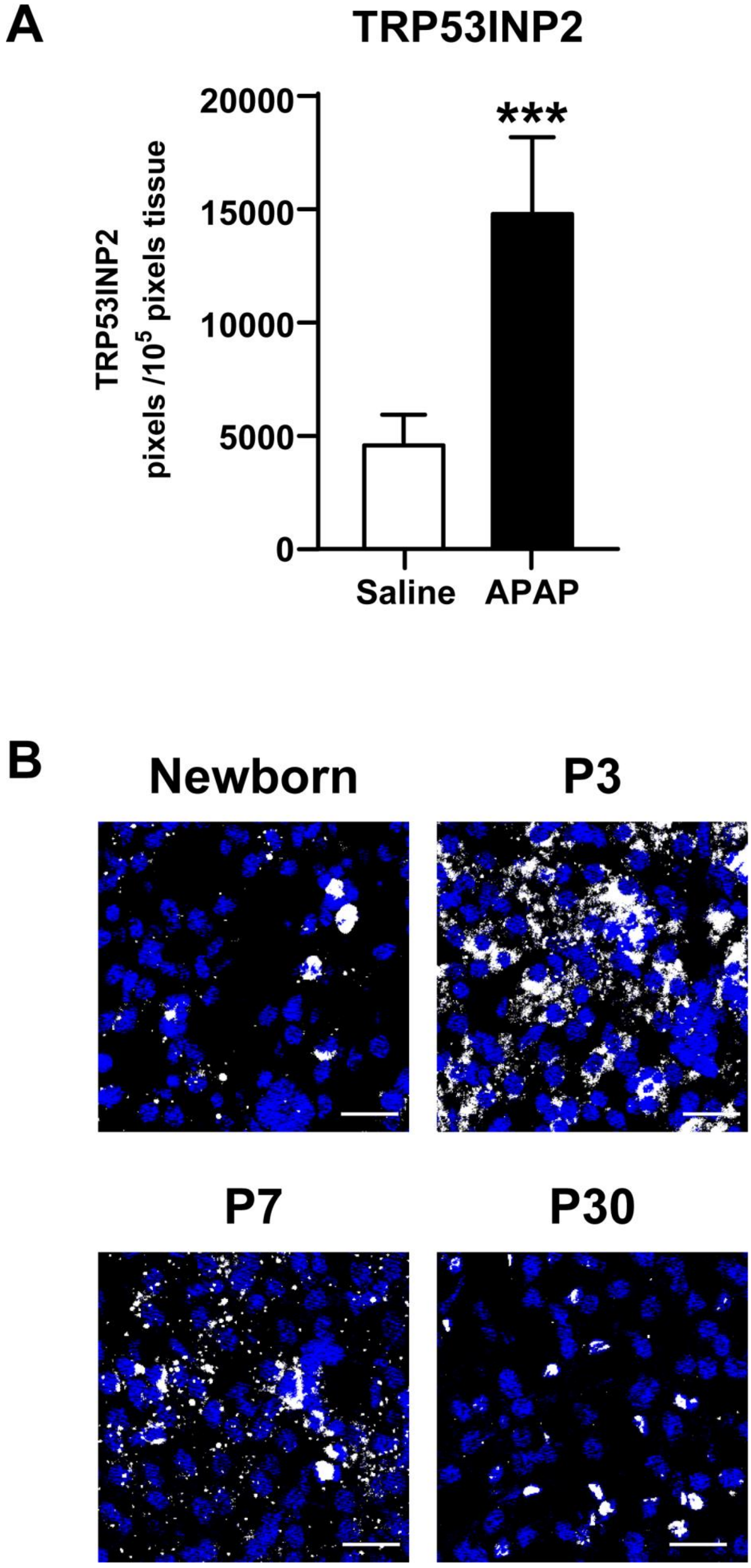

### Supplementary Figure Legends

**Supplementary Figure 1. Development of a canonical TGFβ-signaling reporter mouse model and double TGFβ/BMP reporter mice.** (**A**) The DNA construct used for the generation of TRE-RFPreporter transgenic animals. (**B**) Ex vivo validation of TGFβ responsiveness of the TRE-RFP reporter mouse. Representative images of primary TRE-RFP hepatospheres treated for 48h with the indicated recombinant mouse proteins and/or inhibitors (scale-bars, 50μm). (**C**) Quantitative analysis of RFP levels expressed as Mean Fluorescence Intensity (MFI) in all conditions relative to untreated or diluent-treated hepatospheres, using the ImageJ software. (**D**) Representative images of liver sections derived from the female TRE-RFP/BRE-eGFP reporter animals 72h after intravenous administration of 1x1010 infectious units (i.u.) of Ad0, AdSmad1 or AdSmad3 adenoviruses (scale-bars, 100μm). eGFP (green), RFP (red), Glutamine Synthetase (GS, white), DAPI (blue). (**E**) RT-qPCR analysis of eGFP and RFP mRNA levels in livers of Smad adenovirus-treated animals, relative to Gapdh and normalized to Ad0. Data are expressed as mean±SEM of 3-6 independent samples/condition analyzed using one-way analysis of variance with Bonferroni's post-hoc test

**Supplementary Figure 2. Characterization of eGFP- and RFP-reporter positive cells during liverdevelopment and acetaminophen induced liver injury.** (**A**) Representative confocal images illustrating kinetics of eGFP and/or RFP reporter expression in relation to the establishment of hepatic zone III (scale-bars, 100μm). Liver sections derived from TRE-BRE female reporter animals at indicated developmental stages were stained for eGFP (green), RFP (red), Glutamine

Synthetase (GS, white), and DAPI (blue). (**B**) Representative confocal images showing expression of the eGFP and/or RFP reporter vis-à-vis the expression of GS, 6 and 24h after saline- or APAP-administration. Sections stained for eGFP (green), RFP (red), GS (white) and DAPI (blue). Images in the far right column are magnifications of the areas marked by dashedline squares in the adjacent "merged" images (scale-bars, 100 μm).

**Supplementary Figure 3. Smad7 overexpression in the liver of acetaminophen-treated animals attenuates mobilization of reparatory cells.** Representative confocal images of hepatic tissue sections of PBS- or adenovirus-treated animals, 6 and 24h following APAP or saline administration (scalebars, 100μm), stained for (**A**) αSMA or (**B**) CD45 (white) and DAPI (blue). The lower rows represent magnified areas outlined with dashed-lines in the images directly above. Arrowheads show aSMA positive stellate cells extending from portal towards central regions.

**Supplementary Figure 4. Smad7 overexpression in the liver of acetaminophen-treated animals downregulates the expression of several inflammatory markers.** RT-qPCR analysis of Smad7, Il10, Il13, Il4, and Tnf mRNA levels (versus Cyclophilin A levels) normalized to respective saline mRNA levels. Data are expressed as mean±SEM of 4-5 independent samples/group (male mice) analyzed using one-way analysis of variance with Bonferroni's post-hoc test.

**Supplementary Figure 5. BMP4 treatment of hepatospheres leads to downregulation of senescence-related genes.** Heat-map of the log2 Fold Change values in the ligand-treated hepatospheres, extracted by the KEGG pathway mmu04218 (Cellular senescence).

**Supplementary Figure 6. TRP53INP2 protein is upregulated upon APAP-intoxication and distributed in cytoplasmic puncta during neonatal development.** (**A**) Morphometric analysis for TRP53INP2 immunostaining in Saline- or APAP-treated male C57BL/6 animals, 24h posttreatment. Data are expressed as mean±SEM of 3-4 animals per group, analyzed using unpaired Student's t-test. (**B**) Representative confocal images of hepatic tissue sections isolated from newborn, 3, 7 and 30 days old mice stained for TRP53INP2 (white) and DAPI (blue).
